# Explaining Conformational Diversity in Protein Families through Molecular Motions

**DOI:** 10.1101/2024.02.06.578951

**Authors:** Valentin Lombard, Sergei Grudinin, Elodie Laine

## Abstract

Proteins play a central role in biological processes, and understanding their conformational variability is crucial for unraveling their functional mechanisms. Recent advancements in high-throughput technologies have enhanced our knowledge of protein structures, yet predicting their multiple conformational states and motions remains challenging. This study introduces Dimensionality Analysis for protein Conformational Exploration (DANCE) for a systematic and comprehensive description of protein families conformational variability. DANCE accommodates both experimental and predicted structures. It is suitable for analysing anything from single proteins to superfamilies. Employing it, we clustered all experimentally resolved protein structures available in the Protein Data Bank into conformational collections and characterized them as sets of linear motions. The resource facilitates access and exploitation of the multiple states adopted by a protein and its homologs. Beyond descriptive analysis, we assessed classical dimensionality reduction techniques for sampling unseen states on a representative benchmark. This work improves our understanding of how proteins deform to perform their functions and opens ways to a standardised evaluation of methods designed to sample and generate protein conformations.

## Introduction

Proteins orchestrate all biological processes, and their malfunctions often result in disease. In recent years, high-throughput technologies have greatly improved our knowledge of their amino acid sequences and 3D shapes^1–4^. While reaching the single-structure frontier^5^, these advances have also highlighted the complexities of how proteins move and deform to carry out their biological functions^6,7^. They have stimulated a renewed interest in the modeling of protein and protein complex multiple conformational states^8^. In particular, the success of the protein structure prediction neural network AlphaFold2^9^ has inspired innovative strategies for modifying or repurposing it toward exploring protein conformational space. These approaches involve forced sampling^10^, modulation of input multiple sequence alignment content and depth^11,12^, or guidance with state-annotated templates^13,14^. Although they have achieved promising results for specific protein families, systematic assessments have revealed limitations^15,16^. In addition, studies sampling from low-dimensional representations or manifolds learned from observed or simulated conformations^17–19^ have underscored the difficulty in predicting new, completely unseen states and the importance of high-quality data for training or benchmarking.

Experimental techniques like X-ray crystallography, cryogenic-electron microscopy (cryo-EM), and nuclear magnetic resonance spectroscopy (NMR) are essential for capturing protein functional states^6,20^. The Protein Data Bank (PDB)^4^ offers access to multiple structural states for various proteins, solved independently in different conditions, oligomeric states, and with diverse cofactors and molecular partners. Researchers have actively engaged in efforts to collect, cluster, curate, represent, visualise, and functionally annotate these states^20–23^. These endeavours have provided valuable insights into the biologically meaningful conformational space for specific protein families such as protein kinases^24^, RAS isoforms^25^, ABC (ATP Binding Cassette) transporters^26^, and G-protein coupled receptors (GPCRs)^27^. However, producing or validating functional annotations for structural states involves a substantial amount of manual intervention. Despite the wealth of experimentally resolved protein conformational variability, its full exploitation remains an ongoing challenge.

Ideally, one would like to comprehensively describe protein conformational variability with low-dimensional representations or manifolds amenable to visualisation and interpretation. Principal Component Analysis (PCA) serves as a convenient and robust means to reduce the dimensionality of a dataset, capturing maximum variability^28,29^. The principal components extracted from a conformational ensemble define 3D directions for every atom, and motions along them allow navigating the conformational space^30^. PCA has proven useful for extracting structural transitions from sparse disconnected low-energy structural states^31–36^. Unlike more complex non-linear dimensionality reduction techniques, it offers the advantage of not depending on numerous adjustable parameters and provides a straightforward geometrical interpretation.

Here, we describe a PDB-wide analysis of protein conformational variability across various levels of sequence homology. Our fully-automated computational pipeline, named Dimensionality Analysis for protein Conformational Exploration (DANCE), systematically compiles collections of aligned protein conformations and extracts their principal components. We interpret the representation space defined by the main principal components as the *linear motion manifold* underlying the observed conformations. We provide estimates of the intrinsic dimensionality of these motion manifolds. To assess generative methods, we introduce a benchmark set comprising ten conformational collections representing therapeutic targets with substantial functional transitions. Additionally, we provide baseline performances from classical linear and non-linear manifold learning techniques.

DANCE is versatile, handling both experimental and predicted structures with varying amino acid sequences. It adopts an unbiased approach, avoiding predetermined protein or domain definitions when building the conformational collections. Considering the complete context of input protein chains enables a thorough examination of inter-domain motions. Furthermore, DANCE accommodates uncertainty from unresolved protein regions without assuming potential conformations. It introduces a weighting scheme to mitigate the imbalanced coverage of variables.

We provide several databases of conformational collections representing the whole PDB as well as detailed information about the benchmark on Figshare^37^. In addition, DANCE’s source code is available at: https://github.com/PhyloSofS-Team/DANCE.

## Methods

### Overview of DANCE

DANCE takes as input a set of protein 3D structures (in Crystallographic Information File or CIF format) and outputs a set of protein- or protein family-specific conformational collections or ensembles (in CIF of PDB format). It first clusters and superimposes the input structures based on the similarities found in their corresponding amino acid sequences. The users can choose to analysis all input structures or only those representing monomeric biological units. DANCE then determines the set of principal components sufficient to explain the variability observed within each conformational ensemble. The algorithm unfolds in six main steps depicted in **Fig. 1**.

- **a-Extraction of sequences.** The first step extracts the one-letter amino acid sequences of all polypeptidic chains contained in the input CIF files. In case of multiple models, DANCE retains only the first one. The names of the residues with resolved 3D coordinates are taken from the *_atom_site*.*label_comp_id column*. Residues missing from the protein structure are included as lowercase letters in the sequence if they are defined in the _*entity_poly_seq category*. This information will help in clustering and aligning the sequences (see below). Otherwise, they are replaced by the “X” symbol. The “X” symbol is also used for unknown amino acid types and for modified amino acids without a close natural neighbour. Sequences comprising less than 5 non-“X” residues are then filtered out.
- **b-Clustering of the sequences.** DANCE clusters sequences using MMseqs2^38^. The users can choose the desired levels of sequence similarity and coverage, both set to 80% by default. The coverage is bidirectional by default. This step outputs a TSV file specifying the clusters.
- **c-Multiple sequence alignments.** DANCE then aligns the sequences within each cluster using MAFFT^39^ with default parameters and the BLOSUM62 substitution matrix^40^. It further removes all the columns containing only Xs or gaps, and reorders the sequences according to their PDB codes.
- **d-Extraction of structures.** DANCE extracts 3D coordinates of the backbone atoms N, C, C*α*, and the O atom, of all polypeptidic chains contained in the input CIF files. It reconstructs missing O atoms based on the other atom’s coordinates. It disregards residues with missing backbone atoms and chains shorter than 5 residues.
- **e-Generation of the conformational collections.** DANCE then uses the sequence clusters defined in (b) to group conformations and the residue matching provided by (c) to superimpose them. The superimposition puts their centers of mass to zero and then aims at determining the optimal least-squares rotation matrix minimizing the Root Mean Square Deviation (RMSD) between any conformation and a reference conformation (see below). This is achieved through the ultrafast Quaternion Characteristic Polynomial method^41,42^. The users can choose to account for all the atoms in the superimposition, or only the C*α* atoms. Optionally, the users can filter out the conformations with too few (less than 5 by default) residues aligning to the reference. As a post-processing step, DANCE reduces structural redundancy. Namely, it removes any conformation A deviating by less than *rms*_*cut*_ Å from another one B, provided that the sequence of A is identical to or included in that of B. The value of *rms*_*cut*_ is 0.1 Å by default and is customizable by the users. Finally, DANCE saves the conformational ensemble as a multi-model file in PDB or CIF format. Notice that the models can display different amino acid sequences. DANCE also outputs the corresponding multiple sequence alignments (MSA) in FASTA format, and the matrix of all-to-all pairwise RMSDs.
- **f-Extraction of linear motions.** DANCE performs PCA on the 3D coordinates from each collection. This dimensionality reduction technique identifies orthogonal linear combinations of the variables, namely the Cartesian coordinates, maximally explaining their variance (see below). These linear combinations, which we refer to as principal components or PCA modes, represent directions in the 3D space for every atom. Deforming the protein structure using these components produce motions that connect the conformations observed in the collection. For the sake of simplicity, we directly refer to the principal components as to *linear motions*, although they may not represent actual physical motions undergone by the protein. Furthermore, we estimate the *intrinsic dimensionality* of the linear motion manifold underlying an ensemble’s conformational variability as the number of principal component explaining essentially all its positional variance. The higher the dimensionality – the more complex the linear motions.

**Figure 1.**
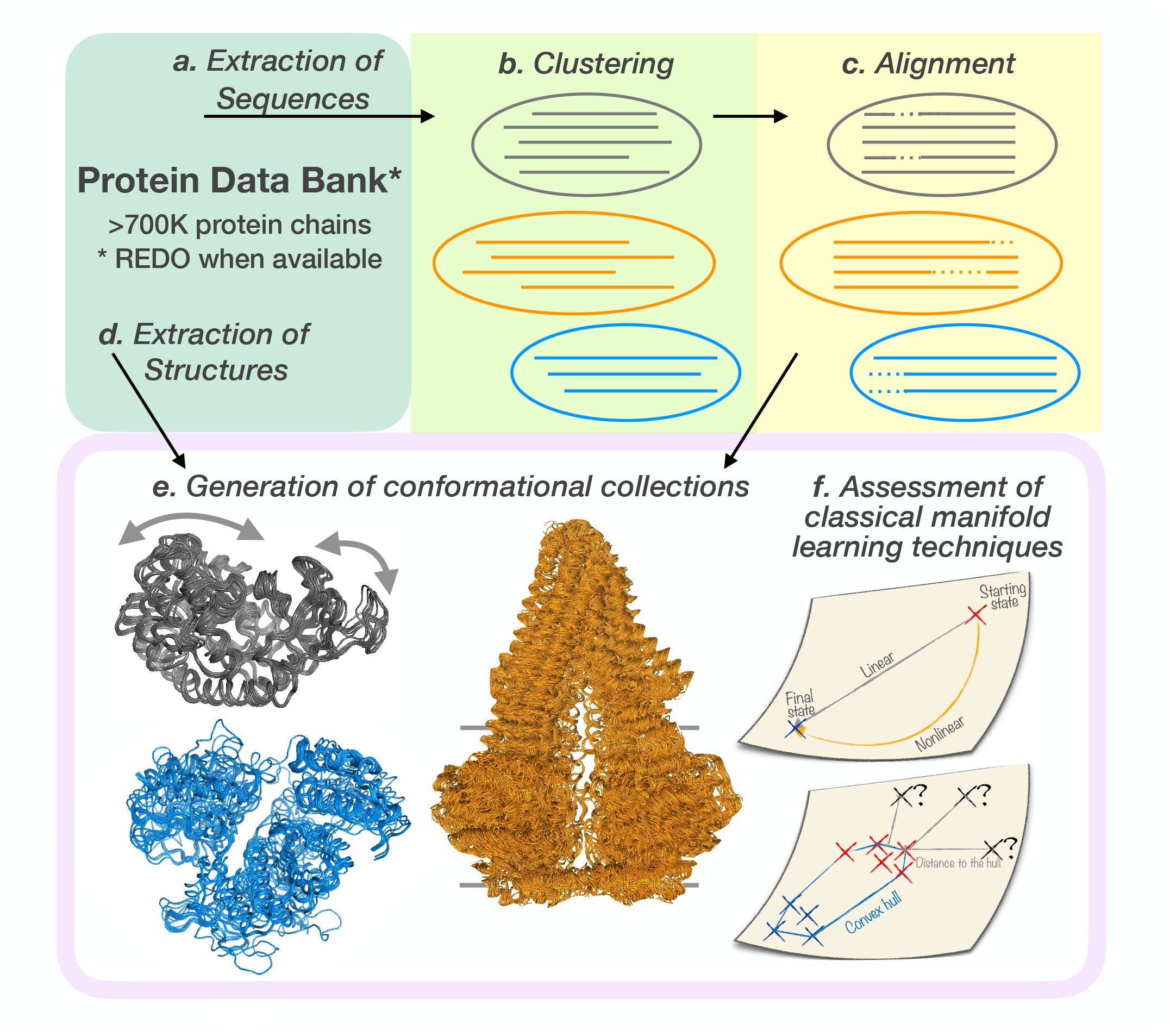
Outline of the study. Our approach, DANCE, exploits both amino acid sequences and 3D coordinates. We applied it to all experimentally determined protein-containing 3D structures from the PDB. Alternatively, users can provide a custom set of experimental structures or predicted models. DANCE first concentrates on sequences. It extracts them from the input structures (A) and clusters them with MMseqs2 based on user-defined similarity and coverage thresholds (B). For each cluster, It generates a multiple sequence alignment using MAFFT (C). It then extracts all 3D coordinates (D), groups the conformations according to the clusters identified in B and superimposes them to generate conformational ensembles (E). The superimposition aims at minimizing the Root Mean Square Deviation to a chosen reference, using the alignments produced by C for mapping the residues. The examples of the bacterial enzymes adenylate kinase (in grey, reference PDB code: 1AKEA) and MurD (in blue, 1E0DA), and the murine ABC transporter P-glycoprotein (5KOYB) are depicted. The arrows indicate adenylate kinase’s main motion. The horizontal lines behind the P-glycoprotein indicate the boundaries for the membrane bilayer. Finally, DANCE summarises conformational diversity through Principal Component Analysis (F). We further assessed the ability of classical manifold learning techniques to reconstruct and extrapolate conformations.

### Choosing a reference

We choose the reference conformation for the superimposition as the one with the amino acid sequence most representative of the MSA. For this, we first determine the consensus sequence s* by identifying the most frequent symbol at each position. We consider “X” symbols as equivalent to gaps. Hence, each position is described by a 21-dimensional vector giving the frequencies of occurrence of the 20 amino acid types and of the gaps. In case of ambiguity, we prefer an amino acid over a gap, hence longer sequences over shorter ones, and an amino acid with a higher BLOSUM62 score over a lower-scored one. Then, we compute a score for each sequence s in the MSA reflecting its similarity to s* and expressed as,

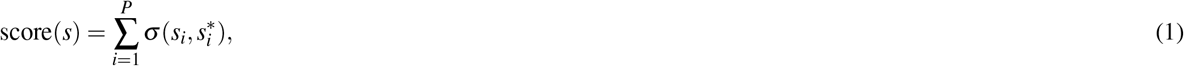

where *P* is the number of positions in the MSA and 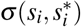 is the BLOSUM62 substitution score between the amino acid *s*_*i*_ at position *i* in sequence *s* and the consensus symbol 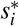 at position *i*. We set the gap score to min_a,b_(σ (a, b)) − 1 = −5.

### Judging the quality of the MSA

We compute the identity level of an MSA as the average percentage of sequence pairs sharing the same amino acid in a column, and the coverage as the percentage of positions having less than 20% of gaps. In addition, we evaluate the global quality of the MSA with a sum-of-pairs score, with σ_*match*_ = 1 and σ_*mismatch*_ = σ_*gap*_ = − 0.5. We normalise the raw sum-of-pairs scores by dividing them by the maximum expected values. The final score for an MSA is thus expressed as,

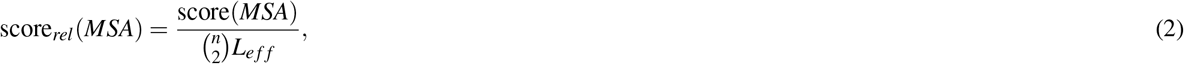

where is the raw MSA score, n is the number of chains or sequences, and *L*_*e f f*_ is the effective length of the MSA, computed as,

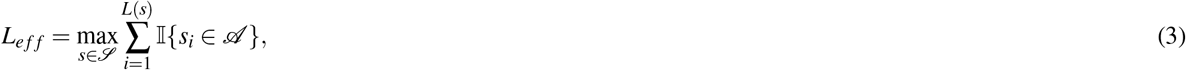

Where ℐ is the set of sequences comprised in the MSA, L(*s*) is the length of the aligned sequence s, and 𝒜 is the 20-letter amino acid alphabet (e.g., excluding gap characters).

### Extracting linear motions

The Cartesian coordinates of each conformational ensemble can be stored in a matrix R of dimension 3*m* × *n*, where m is the number of positions in the associated MSA and n is the number of conformations. Each position is represented by a C-*α* atom. We compute the covariance matrix as,

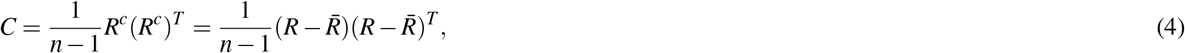

Where 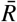 is obtained by averaging the coordinates over the conformations. Alternatively, the users can choose to center the data on the reference conformation. The covariance matrix is a 3*m* × 3*m* square matrix, symmetric and real.

The PCA consists in decomposing *C* as *C* = *VDV*^*T*^ where *V* is a 3*m* × 3*m* matrix where each column defines an eigenvector or a PCA mode that we interpret as a linear motion. *D* is a diagonal matrix containing the eigenvalues. The sum of the eigenvalues 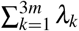 amounts to the total positional variance of the ensemble. The portion of the total variance explained by the *k*th eigenvector or linear motion is estimated as 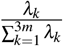.

In addition, we estimate the collectivity^43,44^ of the kth eigenvector as,

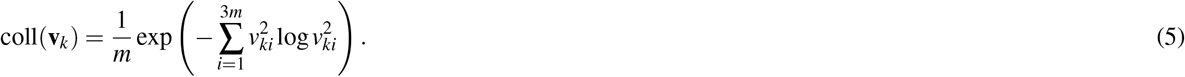

If coll(v_*k*_) = 1, then the corresponding motion is maximally collective and has all the atomic displacements identical. In case of an extremely localised motion, where only one single atom is affected, the collectivity is minimal and equals to 1/*m*.

We also apply PCA to the correlation matrix computed by normalising the covariance matrix as,

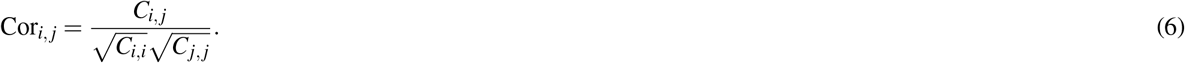

In that case, the sum of the eigenvalues 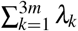 amounts to 1.

Handling missing data

As stated above, the conformations in a collection may have different lengths reflected by the introduction of gaps in the associated MSA. We fill these gaps with the coordinates of the conformation used to center the data (average conformation, by default). In doing so, we avoid introducing biases through reconstruction of the missing coordinates. Moreover, this operation results in low variance for highly gapped positions, thus limiting their contribution to the extracted motions. To go further and explicitly account for data uncertainty, we implemented a weighting scheme. Specifically, DANCE assigns confidence scores to the residues and include them in the structural alignment step and the PCA. The confidence score of a position *I* reflects its coverage in the MSA, 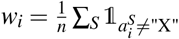, where “X” is the symbol used for gaps. The structural alignment of the *j*th conformation onto the reference conformation amounts to determining the optimal rotation that minimises the following function^45^,

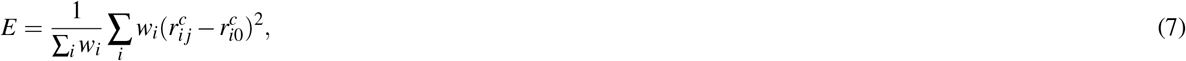

Where 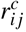 is the *i*th centred coordinate of the *j*th conformation and 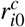 is the ith centred coordinate of the reference conformation. The resulting aligned coordinates are then multiplied by the confidence scores prior to the PCA.

Implementation details

We implemented DANCE in C/C++ and Python. It relies on the C++ GEMMI library^46^ to parse the CIF files and manipulate the structures. It runs MMseqs2 through the following command: *cluster DB clusterDB tmp –cov-mode 0 -c $cov –min-seq-id* $id. It launches MAFFT with the options *auto, amino and preservecase*. The multiple sequence alignment and structure superimposition steps are parallelized. For the PCA, we use the singular value decomposition (SVD) implemented in NumPy^47^ on the R matrix directly. SVD is computationally more advantageous when 3*m* ≫ *n*, which is typically the case of our data, since we only compute the required number of n components. We created structure visualisations in Pymol v2.5.0^48^.

### Application and extension of DANCE

DANCE is applicable to experimental 3D structures as well as predicted 3D models, as long as they comply with the CIF standards.

### Describing conformational variability over the whole PDB

We applied DANCE to all 748 297 protein chains with experimentally resolved 3D structures available in the PDB, as of June 2023. We downloaded all the PDB entries in CIF format from the RCSB^49^. We replaced the raw CIF files with their updated and optimised versions from PDB-REDO whenever possible^50^. It took about 2.25 hours to run DANCE on the whole PDB on a desktop computer with Intel Xeon W-2245 @ 3.90GHz and 32Go of RAM (Supplementary Table S1). The most time consuming steps are the extraction and superimposition of the 3D structures to create the conformational ensembles. We ran DANCE at eight different levels of sequence similarity, designated as 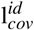, where *id* and *cov* are the sequence identity and coverage thresholds, correspondingly, and range from 50 to 80%. For investigating how the ensembles transformed across levels, we focused on the 18 616 conformational ensembles detected in the most relaxed set up, namely at 30% identity and 50% coverage 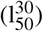. For each ensemble, we extracted its reference protein chain and we traced back the conformational ensembles to which it belonged upon progressively applying stricter thresholds.

### Focusing on the ABC superfamily

We extended DANCE usage beyond the single-chain and sequence-similarity paradigms to describe the conformational variability of ABC (ATP Binding Cassette) transporters. We retrieved a set of 354 ABC protein experimental 3D structures from https://abc3d.hegelab.org^26^. They correspond to functionally relevant states annotated as biological units in the PDB. In most of these structures, several polypeptidic chains, typically 2 or 4, encode the two nucleotide-binding domains (NBDs) and two transmembrane domains (TMDs) of the ABC architecture. In addition, some structures contain several ABC protein copies or some ABC protein cellular partners (small molecules, substrate peptides, interacting proteins). We chose the murine ABC transporter P-glycoprotein (5KOYA) as reference for the subsequent analysis. Its 1182-residue long single polypeptidic chain the full-length transporter architecture.

To cope with the high sequence divergence of the ABC superfamily, we relied on structural similarity for grouping and matching the ABC conformations. Specifically, we used the method Foldseek^51^ to identity structures sharing significant similarity with the reference and align them. We performed a first screen by querying the reference against all individual chains (1 244 in total) and defined significant hits as those with an e-value lower than 10.0. Then, for each structure, we estimated an upper bound on its coverage of the reference by summing up the reference residue ranges appearing in the alignments associated with its significant hits. We filtered out the structures with coverage upper bounds lower than 90%. We performed a second screen by querying the reference against the 209 remaining structures defined as monomers by concatenating their chains. We identified two structures (5NIK, 5NIL) spanning less than 90% of the reference. Permuting their chains did not increase their coverage and thus we removed them. To further detect potentially suboptimal chain orderings, we computed reference to target residue span ratios. We identified one structure, namely 7AHD, with a highly imbalanced ratio of 1.6. Such a high value is indicative of large parts of the reference that could not be aligned to the target structure. Permuting the four chains (A,B,C,D) of 7AHD into (A,D,B,C) led to a more balanced ratio of 0.86. We did not observe discrepancies for other structures and thus we retained their original chain ordering. Finally, we removed the structures with low-quality alignments, i.e., with more than 200 gaps or with a continuous gapped region of more than 60 positions.

Among the 195 structures finally selected, 4F4C, 7SHN and 7AHD contained unknown or unrecognized amino acids which we removed. We ran Foldseek one more time to generate a structure similarity-based multiple sequence alignment centred on the reference 5KOYA. We trimmed the alignment and the 3D structures by removing the residues inserted with respect to the reference. We gave the trimmed alignment and 3D coordinate files as input to DANCE, starting directly from step *d* (see the overview of DANCE algorithm above). For consistency and comparison purposes, we asked DANCE to center the data on the reference. To mitigate the impact of potential alignment errors, we applied weights reflecting position-specific confidence scores (see above, *Handling missing data*). DANCE structural redundancy reduction step removed 7 conformations, resulting in an ensemble of 188 conformations.

We compared this ensemble with those generated by DANCE default sequence similarity-based end-to-end procedure applied to the whole PDB. More specifically, we took the ensembles generated at 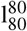 and 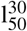 and containing 5KOYA and we rebuilt them with DANCE, applying the 5KOYA centering and the uncertainty weighting scheme. We estimated the similarity between the ensembles’ motion subspaces as the Root Mean Square Inner Product (RMSIP)^52,53^. The latter measures the overlap between all pairs of the l first PCA modes and is defined as,

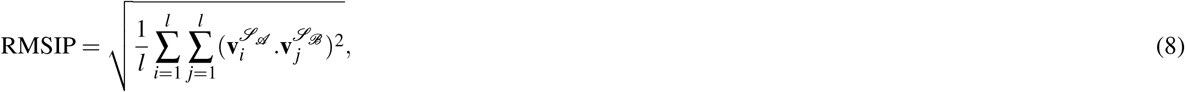

Where 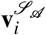 and 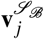 are the *i*th and *j*th PCA modes extracted from the conformational ensembles ℒ_𝒜_ and ℒ_ℬ_, and l is the number of modes considered for the comparison. Moreover, we monitored the distance between the geometric centres of the two NBDs defined by the C-*α* atoms of residues numbered 346-596 and 929-1182, respectively, in the reference 5KOYA.

### Benchmarking for the generation of unseen conformations

We further investigated whether the extracted linear principal components could be useful to predict unseen conformations. Moreover, since the manifold underlying our data is a *priori* non-linear, we tested whether non-linear methods could achieve better reconstructions than linear PCA. We focused on the widely used kernel Principal Component Analysis (kPCA)^54,55^ and the uniform manifold approximation and projection (UMAP)^56^.

### Dimension reduction with non-linear kernel PCA

The intuition behind kPCA is to map the input data points to a higher dimensional space where they will be linearly separable by a classical PCA. The mapping function *ϕ* : ℝ^3*m*^ → ℝ^*M*^ is not known. Instead of explicitly calculating it, we use a kernel function *k*(***r***_*i*_, ***r***_*j*_) = *ϕ* (***r***_*i*_)^*T*^*ϕ* (**r**_*j*_), where ***r***_*i*_ and ***r***_*j*_ are two conformations. We considered three commonly used kernels,

- the polynomial kernel 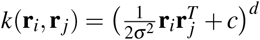, where *c* = 1 and *d* = 3 by default,
- the sigmoid kernel 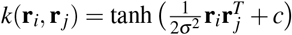, where *c* = 1 by default,
- and the radial basis function (RBF) or Gaussian kernel 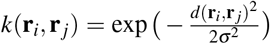, where *d*(**r**_*i*_,***r***_*j*_) is the Euclidean distance between the two conformations **r**_*i*_ and ***r***_*j*_.

We explored different values of the hyperparameter σ. For sufficiently large values, i.e., 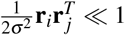 or 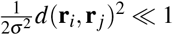 the kernel becomes effectively linear.

Thus, given the input coordinates *R* representing *n* conformations, we computed the corresponding kernel matrix *K* of dimension *n* × *n* and decomposed it using the classical PCA. The resulting principal components {*v*_**1**_, *v*_**2**_, …, *v*_**n**_} can then be expressed as,

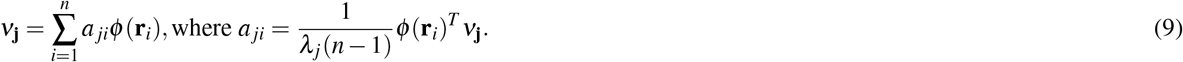

### Uniform manifold approximation and projection

The UMAP algorithm first builds a graph representing the data in the ambient space, and then determines the most similar graph in a lower dimension. It relies on the assumptions that there exists a low-dimensional manifold on which the original data would be uniformly distributed and that this manifold is locally connected. Under such assumptions, any ball of fixed volume on the low-dimensional manifold should contain approximately the same number of points. Thus, to build the graph, UMAP defines balls in the ambient space centred at each point and encompassing its *n*_*neigh*_ nearest neighbours. The balls have variable sizes that reflect the topology of the dataset in the ambient space. UMAP then connects points whose corresponding balls overlap and computes the edge weights by combining the balls’ radii. The resulting graphical representation is projected into a lower-dimensional space by minimising the cross entropy between the high- and low-dimensional graphs, which can be viewed as a force-directed graph layout algorithm. We explored two hyperparameters, namely the number of neighbours *n*_*neigh*_ controlling the balls’ radii and the minimum distance *d*_*min*_ apart that points are allowed to be in the low dimensional representation. Low values of *n*_*neigh*_ will make UMAP focus on local details of the dataset topology while high values will account for more global properties. Increasing *d*_*min*_ will push points far from each other in the representation space.

### Generating conformations

For linear PCA, generating 3D conformations by combining the principal components is straightforward. More specifically, given a set of l PCA modes computed from the coordinates *R*, we generate a new conformation 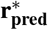 as,

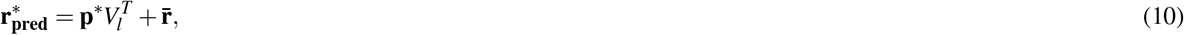

where the matrix V_*k*_ ∈ ℝ^*3m×l*^ contains the modes, 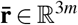 is the average conformation, and **p**\* ∈ ℝ^l^ is a point in the *l*-dimensional representation space defined by the modes. The coordinates of **p**\* specify the amplitudes of the modes.

For kPCA and UMAP, we need to learn an inverse transform function that maps points in the *l*-dimensional representation space defined by the components back to the input space. This problem is known as the *pre-image problem*. To solve it for kPCA, we used kernel ridge regression of the input coordinates R on their low-dimensional projections in the representation space as described in^57,58^ and implemented in the scikit-learn Python library^59^. The contribution of the L2-norm regularisation is controlled through the hyperparameter *α*. More technically, *α* connects the squared L2-norm between a point in the representation space and its reconstruction with the squared L2-norm of the kernel weights used for the reconstruction. In the case of UMAP, we used the built-in inverse_transform function^60^. It relies on stochastic gradient descent to minimise the cross entropy between the low-dimensional graph and its high-dimensional pre-image graph.

### Leave-one-cluster-out cross-validation procedure

We assessed the predictive performance of PCA and kPCA with a *leave-one-out* cross-validation procedure. Since the conformations are not evenly distributed within an ensemble, we grouped them into clusters prior to the evaluation. We performed the clustering in the l-dimensional PCA representation space, where *l* is the minimal number of linear components sufficient to explain 90% of the ensemble’s total positional variance. We used the *k*-means clustering^61^ with *k* = *l* + 2.

Given a clustered ensemble, we systematically tested the ability of the principal modes inferred from *l* + 1 clusters to predict the conformations belonging to the held-out cluster. We reconstructed each test conformation **r**\* from its projection **p**\* in the l-dimensional representation space. For the classical PCA, we computed the projection as,

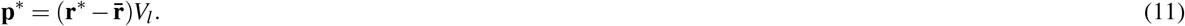

For the kPCA, the projection onto the principal component *v*_j_ is expressed as,

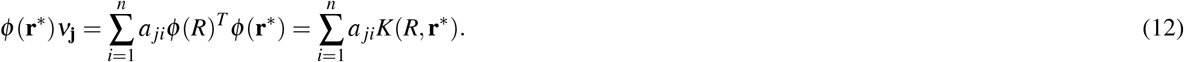

We evaluated the reconstruction error as the RMSD between the predicted conformation 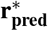 and the original conformation **r**\*.

### Distance to the training set

We estimated the difficulty of reconstructing a given conformation by computing its distance to the convex hull defined by the conformations used for training in the *l*-dimensional representation space. Setting the number of clusters in the training set to *l* + 1 ensures that the convex hull will be a polytope of dimension at least *l*. For instance, in 1 dimension, we need at least 2 affine-independent points to define a 1-polytope. The explicit computation of the convex hull of n points in l dimensions is an operation whose complexity is of the order of *O*(*n*^*l*/2^)^62^ and rapidly becomes computationally infeasible as the value of *l* increases. Nevertheless, the calculation of the distance of a given point to the hull does not require computing the convex hull explicitly and is a much simpler computational problem. It can be solved in quasilinear time with quadratic programming (QP). Here, we used the efficient and exact QP simplex solver proposed in^63^ and implemented in the Computational Geometry Algorithms Library (CGAL)^64^. It takes advantage of the low dimensionality of the representation space by observing that the closest features of two *l*-polytopes are always determined by at most *l* + 2 points.

In order to compare distances across systems of different sizes, we scale them by the number of positions *m*,

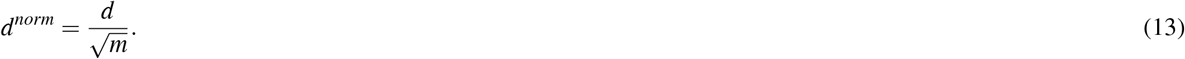

This normalisation also allows relating distances in the representation space with RMS deviations in the 3D Cartesian space. Indeed, let us consider an ensemble of conformations exhibiting a purely one-dimensional motion. Any two conformations distant by an RMSD of 1 Å in the original 3D space will be separated by a normalised distance of 1 Å in the one-dimensional representation space.

### Interpolating between states

We generated interpolation trajectories between ATPase states with PCA and kPCA. We started from the conformational clusters defined in the leave-one-out procedure and identified clusters 0 and 4 as the most extreme ones along the first PCA component. Secondly, we used these two clusters only to learn PCA and kPCA low-dimensional representation spaces. We computed the coordinates of the clusters’ centres in these spaces and defined interpolation trajectories between them with 50 regularly spaced intermediate points. We then generated 50 conformations from the 50 intermediate points. We finally determined the minimal RMS deviation between each generated conformation and the known conformations from clusters 1, 2 and 3. We qualitatively compared these trajectories with physics-based non-linear trajectories computed with NOLB^90^. NOLB extracts normal modes from a starting conformation and models the transition to a target conformation as a series of twists extrapolated from these modes with optimal amplitudes, as described in^91^. We chose 1KJUA from cluster 0 as the starting conformation and 1T5SA from cluster 4 as the target conformation.

## Results

We used DANCE to chart the experimentally resolved conformational diversity of protein families (**Fig. 1**). We explored eight levels of sequence similarity (sim) and coverage (cov), denoted as 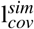, to group the ∼750K chains included in the PDB as of June 2023 (**Supplementary Fig. S1A** and **Supplementary Table S2**). In the most conservative set up, namely 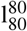, less than 3% of the conformations remain isolated (**Supplementary Fig. S1A**, singletons). Most of the conformational collections (or ensembles) are associated with multiple sequence alignments of high quality across all levels (**Supplementary Fig. S1B**). Sequence identity and coverage are more widely distributed in more relaxed conditions, but the median values always remain very high, above 0.95 (**Supplementary Fig. S1C-D**).

### Experimentally resolved conformations lie on low-dimensional manifolds

Only one or two linear principal components suffice to explain almost half of the ensembles’ conformational diversity (**Fig. 2a**). We interpret these components as directions of motion, and by simplification, we will denote them as linear motions in the following (*see Methods*). In the overwhelming majority of cases, less than eight linear motions explain more than 90% of the total positional variance. These observations hold true across all sequence identity and coverage levels. They indicate that the conformational states captured by experimental techniques for a protein or a protein family lie on a low-dimensional manifold. This low dimensionality is only partially determined by the cardinality of the ensembles (**Supplementary Fig. S2A-B**). Almost 30% of the most highly populated ensembles (>50 conformations) detected at 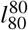 can be comprehensively described with less than three linear motions (**Supplementary Fig. S2C**). This proportion increases up to 46% in the most relaxed conditions, namely at 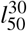 (**Supplementary Fig. S2D**).

**Figure 2.**
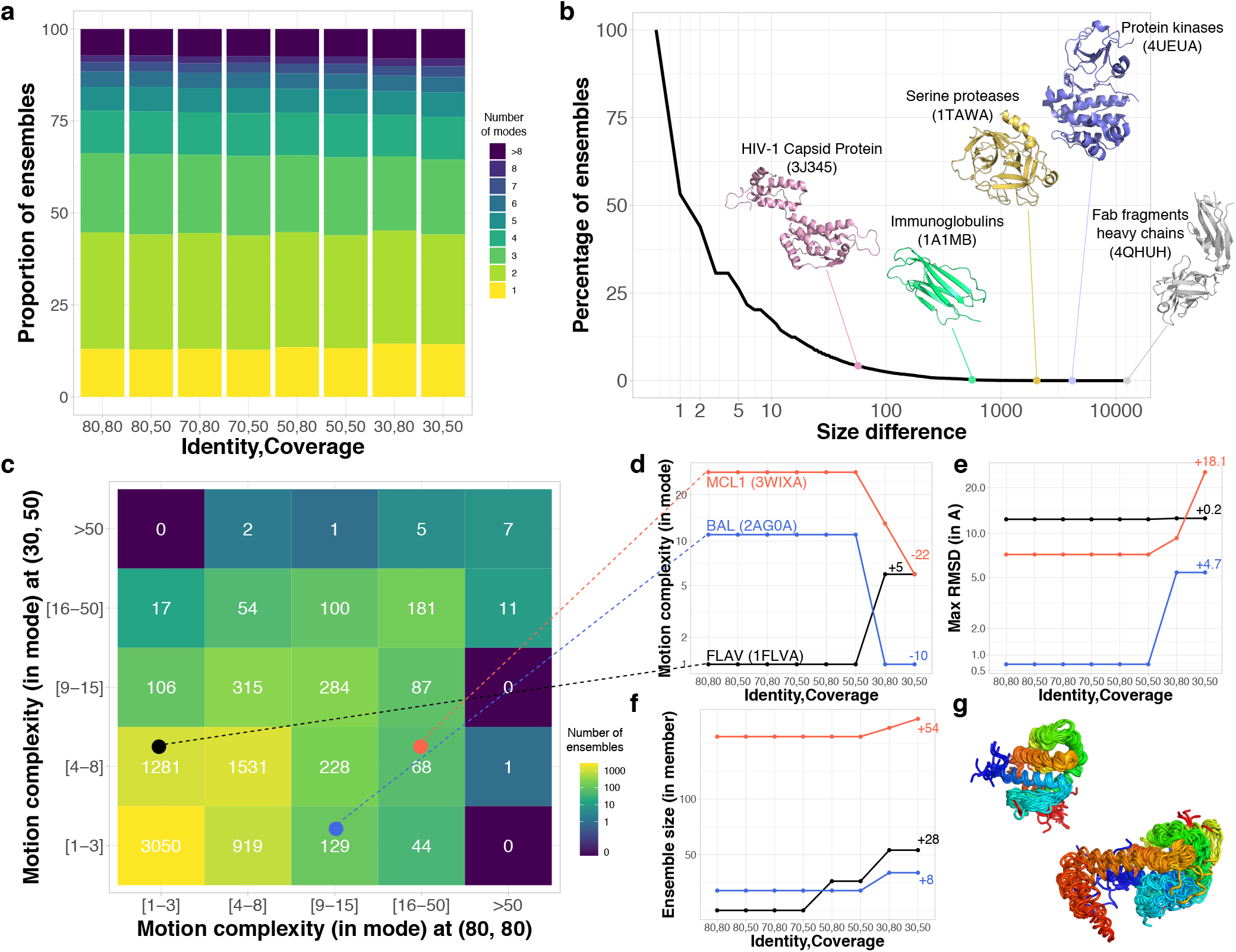
Evolution of protein conformational diversity across sequence similarity levels. **a**. Proportion of conformational ensembles requiring n linear PCA modes to explain 90% of their total positional variance, with n varying from 1 to 8. The number of modes n is an indicator of motion complexity. Singletons and pairs are excluded. **b**. Cumulative distribution of the number of conformations gained from the most stringent level, namely 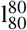, with 80% sequence similarity and coverage, to the most permissive one, 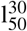, with 30% similarity and 50% coverage. The 3D structures of the reference protein chains are depicted for a few ensembles. **c**. Comparison of motion complexity between the most stringent and most relaxed set ups. We considered only the cases where the ensemble at 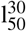 is bigger than the corresponding one at 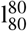. Singletons and pairs are excluded. D-G. Detailed evolution of three ensembles marked by colored dots in panel C. **d**. Motion complexity expressed as a number of modes. The names and PDB codes of the reference chains are indicated. **e**. Motion amplitude, measured as the maximum RMSD between any two conformations (in Å). **f**. Conformational collection size. **G**. Conformational diversity observed for the Bcl-2 family. On the top left, the 54 conformations comprised in the MCL1 ensemble at 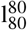. At the bottom right, the 218 additional conformations at 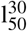. The color code indicates the position in the sequence, from the N-terminus in blue to the C-terminus in red.

The bacterial adenylate kinase gives an example of a one-dimensional motion underlying its 42 conformations (**Fig. 1e,** in grey). One can easily classify the conformations by visual inspection into two main states, open and closed, deviating by about 7 Å. The bacterial enzyme MurD (Fig. 1e, in blue) and the murine ABC transporter P-glycoprotein (**Fig. 1e,** in orange) also exhibit low-dimensional opening-closing motions. In particular, the P-glycoprotein’s collection reveals a rich spectrum of intermediate conformations between the open and closed forms (**Fig. 1e,** in orange). The main motion involves about 70% of the protein and modulates the volume of the transporter’s internal cavity within the lipid bilayer up to over 6,000 Å^3^^65^. It explains about 80% of the total positional variance on its own. The remaining variability is mostly due to rotations of the nucleotide binding domains with respect to the transmembrane helical bundles and to loop deformations.

### A few protein families display huge conformational expansion upon relaxing the sequence selection crite-ria

To investigate how the conformational ensembles transformed with sequence similarity, we systematically backtracked the 18 616 representative protein chains identified at 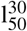 across more stringent levels (*see Methods*). The fragment antigen-binding regions display the largest growth between the most stringent and most relaxed sequence selection criteria (**Fig. 2**). For instance, while the Fab6785 light chain’s ensemble at 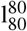 comprises a bit less than 300 conformations, it expands up to over 12 500 conformations at 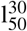 (**Fig. 2b**, PDB id: 4QHUH). With the largest number of conformations at 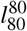, the HIV-1 capsid protein’s ensemble however displays a relatively limited expansion across the different levels, from 3 334 to 3 391 (**Fig. 2b**, 3J345). Bovine trypsin and its close homologs give an example of an extensively characterized subfamily, with 470 different conformations detected at 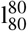. This ensemble expands by more than 5 folds, aggregating different serine proteases, upon relaxing the criteria to 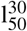 (**Fig. 2b**, PDB id: 1TAWA). Likewise, the Beta-2-microglobulin and its close homologs have a large body of 1 465 conformations at 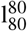, growing further up to 2 025 conformations at 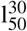 by including other immunoglobulins (**Fig. 2b**, 7MX4B). By contrast, the reconstructed ancestral tyrosine kinase AS, a common ancestor of Src and Abl, has only 2 conformations available in the PDB and no close homologs. At 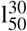, it serves as representative for a huge ensemble of over 4 000 protein kinase conformations (**Fig. 2b**, 4UEUA). Apart from these over-represented protein families or superfamilies, the ensembles generally gain only a few conformations, with a median value of 4.

### Family expansion may lead to an apparent motion simplification

As an ensemble grows, the gained conformations may lie on the same motion manifold, defined by the subset of principal components explaining the variance, or give rise to new motions represented by new components (**Fig. 2c**). The bacterial long-chain flavodoxin exemplifies the second scenario (**Fig. 2d-f**, in black). At 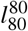, it undergoes a one-dimensional motion describing the transition between a compact state and a partially unfolded conformation (**Supplementary Fig. S3**). Upon relaxing sequence similarity to 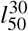, the ensemble roughly doubles in size (**Fig. 2f**) and the newly added conformations exhibit complex deformations of the FMN binding pocket. As a result, five more linear motions are required to explain the positional variance (**Fig. 2d**). Hence, in this case, the motions get more complex when considering more distant homologs.

The emergence of new motions does not however systematically lead to an increased motion complexity. The murine MCL1 gives an illustrative example of apparent motion simplification upon expansion (**Fig. 2d-f**, in red, and **Fig. 2g**). At 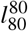, almost 30 components are needed to explain the variability observed over the couple of hundreds conformations in the ensemble. They represent local deformations of the inter-helical loops and the extremities (**Fig. 2g** and Supplementary **Fig. S3**). Extending the ensemble to distant members of the Bcl-2 family brings in about 50 new conformations (**Fig. 2f**). They reveal a new extended state the protein BAX adopts upon assembling into domain-swapped dimers^66^. The large amplitude transition between the compact conformation and the extended one takes a big part in the variance, resulting in a drastically reduced motion complexity (**Fig. 2d**). The benzaldehyde lyase BAL gives another example (**Fig. 2d-f**, in blue) where the transition to a new state, adopted by the distant homolog actinobacterial 2-hydroxyacyl-CoA lyase^67^, dominates the variance (**Supplementary Fig. S3**). The conformational variability transforms from small (<1Å) seemingly random fluctuations to a one-dimensional motion.

Overall, about a third of the ensembles undergo an apparent motion simplification upon expansion (**Fig. 2c** and **Supplementary Fig. S4A**). They likely represent protein families where distant homologs exhibit novel distinct states. The larger the deviations of these novel states with respect to the other ones, the higher the contribution of the corresponding motions to the variance. To mitigate this variance-dependent effect, we repeated the analysis on the correlation matrix. The latter estimates the extent to which the residues move in the same direction, regardless of the magnitude of their displacements. We found that the motion complexity still decreases in over 20% of the ensembles (**Supplementary Fig. S4B**). This result indicates that motion simplification does not merely reflect larger transitions “hiding” smaller rearrangements. A substantial fraction of protein families show evidence of more concerted residue movements between more distant homologs.

### Beyond single chains and sequence similarity, the ABC superfamily as a case study

We explored the possibility of using DANCE to chart the conformational variability of remote homologs with low sequence similarity and variable chain composition. We focused on the ABC (ATP Binding Cassette) transporter superfamily. The ABC architecture comprises two nucleotide-binding domains (NBDs) and two transmembrane domains (TMDs) encoded by one or several polypeptidic chains (**Fig. 3a**). The NBDs are highly conserved across species and families, whereas the TMDs exhibit various scaffolds associated with heterogeneous transport functions^26^. We considered a collection of a few hundreds ABC protein experimental 3D structures^26^, taking the single-chain murine P-glycoprotein as reference (**Fig. 3a**, 5KOYA).

**Figure 3.**
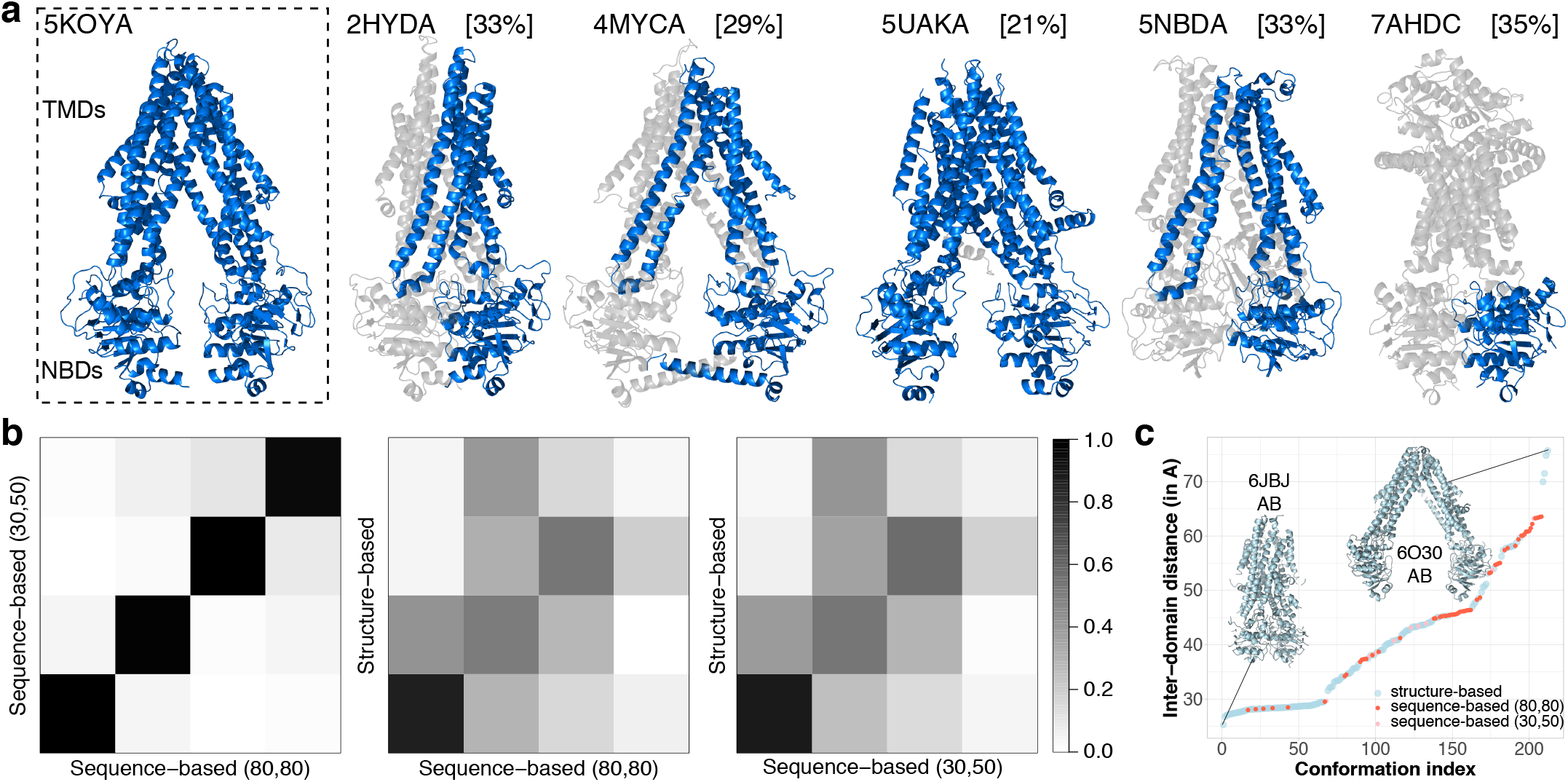
ABC transporters’ conformational variability. **a**. Examples of protein structures from the ABC structure similarity-based conformational collection. The reference chain (5KOYA) is on the left, where we indicate the location of the two NBDs (∼ 500 residues) and two TMDs (∼700 residues). Within each of the other structures, we highlight one chain in marine, give its percentage of identity with the reference in squared brackets, and display the remaining chains in transparent grey. The six marine chains were assigned to six different collections by DANCE’s default sequence similarity-based end-to-end protocol at 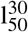. **b**. Comparison of motion subspaces extracted from the sequence-based ensembles at 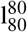 (61 conformations) and 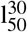 (71 conformations) and the structure-based one (188 conformations). Each matrix shows the absolute pairwise scalar products computed for the first four PCA modes. The corresponding RMSIP are 0.99, 0.71 and 0.73. **c**. Distance between the geometric centres of the two NBDs (in Å). The conformations are ordered along the *x*-axis from the most closed one to the most open one.

We bypassed DANCE sequence extraction, clustering and alignment steps and directly gave it a pre-computed alignment built from structural similarities as input (*see Methods*). Relying on structure rather than sequence similarity and considering various oligomeric states provided a more comprehensive description of ABC transporters’ functional motions and states (**Fig. 3** and **Movies S1-2**). The resulting ensemble comprises 188 conformations encompassing 295 protein chains, some of which have sequence identity below 30% or coverage lower than 50% (**Fig. 3a**). A set of 25 linear motions are required to explain the positional variance. By comparison, the sequence similarity-based 5KOYA-containing collection generated by DANCE at 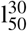 contains only 71 conformations explained by only four linear motions. These motions are essentially identical to those extracted from the 61 conformations at 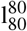 (**Fig. 3b**, RMSIP = 0.99).

Despite having different motion complexities, the sequence- and structure-based conformational collections have largely overlapping motion subspaces (**Fig. 3b**, RMSIP ∼ 0.7). In particular, they all share the same most contributing motion describing the transition between the transporter inward-closed and inward-open forms (**Supplementary Fig. S5**). This functional transition controls the substrate access to the transporter’s central binding pocket. It explains 45 to 70% of the variance on its own and involves over two-thirds of the residues. The structure similarity-based collection represents a quasi-continuum of increasingly open states (**Fig. 3c**, in blue, and **Movie S1**) between two extreme dimeric forms, one from the human lysosomal cobalamin exporter ABCD4 where the two NBDs are in contact and the other from Salmonella typhimurium’s lipid A transporter MsbA with a widely open cavity. The overwhelming majority of conformations are regularly spaced by inter-NBD distance increments smaller than 1 Å. By contrast, the sequence similarity-based collections populate sparse regions of this continuous transition, with a high concentration of semi-open and open states (**Fig. 3c**, in pink and red, and **Movie S2**).

### Classical manifold learning techniques can generate highly accurate conformations

Beyond describing the observed conformational variability, we evaluated the ability of several popular manifold learning techniques to generate unseen conformations. To do so, we identified a set of ten conformational ensembles with very different degrees of motion complexity (**Fig. 4a** and **Supplementary Table S3**). They comprise between 20 and over 3 300 conformations and their reference chains contain 80 to 1 200 residues. They represent proteins or protein families displaying substantial (≥ 5 Å) and functionally relevant conformational changes, namely adenylate kinase (ADK)^68,69^, MurD^19,70^, the calcium pump ATPase^71,72^, the ABC transporters^26,73^, the small heat shock protein *α*B crystallin (Crys)^74,75^, the heat shock protein HSP90^76,77^, calmodulin (CALM)^78,79^, kinases (KIN)^80,81^, RAS^25,82^, and the HIV capsid protein (CAP)^83,84^. Most of them have been extensively characterized by experimental structure determination techniques or computational methods for simulating protein dynamics. Targeting their motions or their specific conformations bears a therapeutic interest.

**Figure 4.**
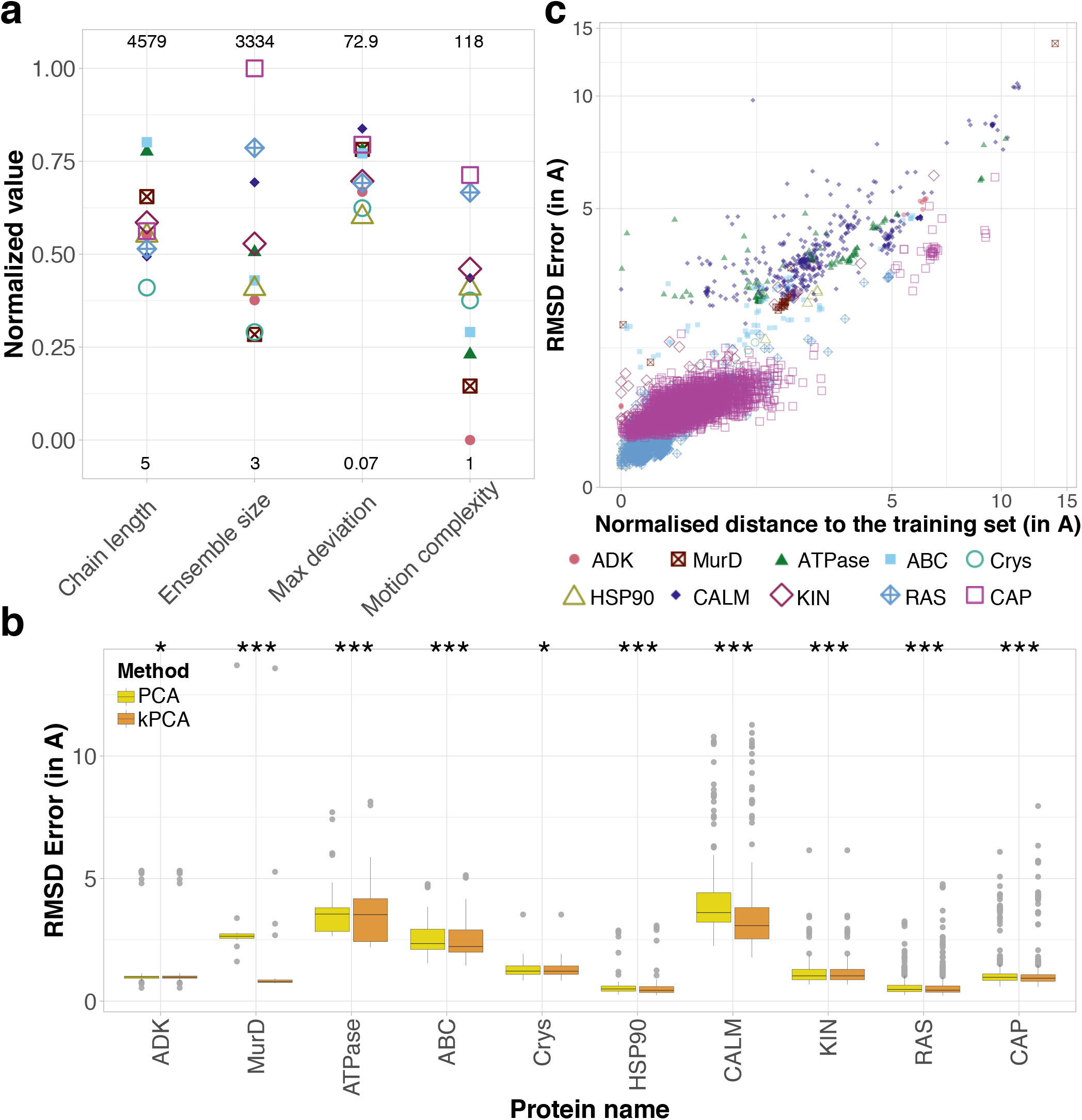
Assessment of classical manifold learning techniques. **a**. Properties of the benchmark set. For each property y, we computed its normalised value as 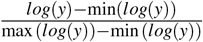. The minimum and maximum are determined over the whole 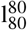 database. They are given at the bottom and on the top, respectively. **b**. Distributions of the RMSD reconstruction errors (in Å) for each ensemble in the benchmark set. We systematically reconstructed each conformation through a leave-one-cluster-out cross-validation procedure (see Methods). We set the two hyper-parameters of the kPCA (RBF kernel) to the values yielding the best reconstruction, for each ensemble. The protein names in the x-axis are ordered according to motion complexity. The stars indicate the statistical significance of the better performance of kPCA compared to linear PCA (one-sided paired t-test; *: p-val < 1*e*^−2; ***^: p-val< 1*e*^−5^). c. RMSD reconstruction error in function of the distance to the training set’s convex hull in the PCA representation space.

We chose the linear PCA as baseline and we considered four non-linear techniques, namely kernel PCA (kPCA)^54,55^, UMAP^56^, isoMAP^85^ and t-SNE^86^. While all techniques allow for projecting the conformations in a low-dimensional space, only PCA, kPCA and UMAP allow for reconstructing conformations from the projections through an inverse transform. Furthermore, UMAP is limited to a narrow range of dimensions and, as a consequence, we could apply it only to a subset of the benchmark (see Methods). Hence, we primarily focus on the comparison between PCA and kPCA in the following. We tested three different kernels for kPCA, namely the sigmoid, polynomial and radial basis function (RBF) kernels. Within each ensemble, we first learned low-dimensional representations of a subset of conformations used as training samples. We then projected the test conformations, not seen during training, to the learned representation space, and mapped the projections back to the original 3D Cartesian space. The mapping is determined analytically in the case of linear PCA and learned in the case of kPCA and UMAP (*see Methods*). We evaluated the quality of the 3D reconstructions by computing their RMS deviations from the original conformations. We found that both PCA and kPCA (with RBF kernel) produced high-accuracy reconstructions (RMSD error below 2Å) for almost all proteins (**Fig. 4b**). The error distribution median and width vary from one protein to another and do not depend on motion complexity. For instance, all reconstructed conformations of HSP90 deviate by less than 3 Å from the original ones, while the reconstruction error can be as high as 14 Å for MurD. The distributions are overall shifted toward higher reconstruction errors for ATPase and ABC, likely due to their large size (∼1 000 amino acids compared to less than 500 for the other proteins, **Supplementary Table S3**), and for CALM, likely due to the large amplitude of its motions (average RMSD = 10.38 ± 4.23 Å, **Supplementary Table S3**). The nonlinear kPCA performed significantly better than the linear PCA for all proteins from the benchmark. It allows increasing the percentage of high-quality reconstructions (RMSD error<2Å) from 5 to 82% for MurD and from 18 to 26% for ABC (**Supplementary Table S4**). Nevertheless, the reconstruction accuracy of kPCA varies greatly depending on the values of the two hyperparameters controlling the kernel width and the amount of regularisation (**Supplementary Fig. S6**). The optimal values vary from one system to another and determining them *a priori* is not trivial. The sigmoid and polynomial kernels may be better suited than RBF for some of the proteins, but the results are overall similar (**Supplementary Fig. S7** and **Supplementary Table S5**). By contrast, UMAP consistently produced reconstructions of substantially lower accuracy than PCA and kPCA (**Supplementary Fig. S7** and **Supplementary Table S5**). Moreover, its runtime was 100 to 100K times longer, depending on the representation space dimension.

### Reconstruction accuracy strongly depends on the distance to the training set

The quality of the predictions strongly correlates with the distance between the test conformation and the training set’s convex hull in the low-dimensional representation space (**Fig. 4c**). The linear PCA produces highly accurate reconstructions, with an RMSD error smaller than 2 Å, only for conformations lying in a close vicinity to the training set’s convex hull (distance smaller than 3 Å). We observed a similar tendency for kPCA (**Supplementary Fig. S8**). This dependence can be appreciated by visualising how the conformations cluster in the representation space (**Supplementary Fig. S9**). For instance, the most poorly reconstructed MurD conformation forms a singleton located far away from all other conformations, particularly along the first most important principal component (**Supplementary Fig. S9B**, dark dot). For this protein, the kPCA performed substantially better than the PCA thanks to a better reconstruction of the most populated cluster (**Supplementary Fig. S9B**, light squares). Hence, the further away from the training set, the more difficult the task. In addition, the overwhelming majority of conformations lie outside of the training set’s convex hull. This observation agrees with a recent study showing that interpolation almost surely never happens with high dimensional datasets^87^. The 14 conformations (out of 4 892 in total) located inside come from ADK, CALM, KIN, RAS and CAP and are all reconstructed with high accuracy, the RMSD errors ranging from 0.2 to 2.9 Å.

### Stereochemical quality and biological significance of the generated conformations

We assessed the physical realism of the generated conformations with PROCHECK, a popular software for checking the stereochemical quality of protein conformations, by comparing them with expected statistics^88^. The PCA- and kPCA-generated conformations displayed proportions of residues in the most favoured (or core) regions of the Ramachandran plot comparable with the experimental conformations (**Supplementary Fig. S10**). In particular, most of the conformations generated by kPCA for ADK, MurD, Crys, HSP90, RAS and CAP had more than 90% of their residues in the most favoured regions. Some of the generated conformations were even of higher stereochemical quality than their experimental counterparts. For instance, for the protein RAS, the linear PCA reconstruction greatly improved over the crystallographic structure 1PLL (chain A), from 63.6% to 94.4% residues in the most favoured Ramachandran regions. The secondary structures in the generated conformation are visibly better defined than in the experimental one (**Supplementary Fig. S11**). In this case, the PCA was able to denoise a poorly resolved conformation by learning from the other conformations in the collection. The conformations generated for CALM have the lowest stereochemical quality (**Supplementary Fig. S10**), in line with their large RMSD errors (**Fig. 4b**). The conformations generated with UMAP have very poor quality across all proteins to which we applied it (**Supplementary Fig. S10**, in green blue).

We further probed the biological significance of the representation spaces learnt by PCA and kPCA by investigating whether linear interpolations between extreme states in these spaces could recapitulate known intermediate conformations. We focused on ATPase as a case study and we chose the centres of clusters 0 and 4 as the end points (**Supplementary Fig. S9C**). We first learnt a low-dimensional representation space using all conformations from the two clusters, and we then generated 50 regularly spaced intermediate conformations along the trajectory between them. The generated conformations approximate known intermediates with RMSD errors as low as 3.6Å in the first half of the trajectory and 3.8Å in the second half (**Fig. 5a**). These results suggest that interpolating between known states in the learnt representation space can be a valid strategy to generate plausible intermediate conformations. In addition, one can visually appreciate the non-linear nature of the trajectories computed with kPCA compared to the linear PCA (**Fig. 5b**, compared left and middle panels). They bear some resemblance with trajectories computed using non-linear normal mode analysis^89–91^ (**Fig. 5b**, compared middle and right panels).

**Figure 5.**
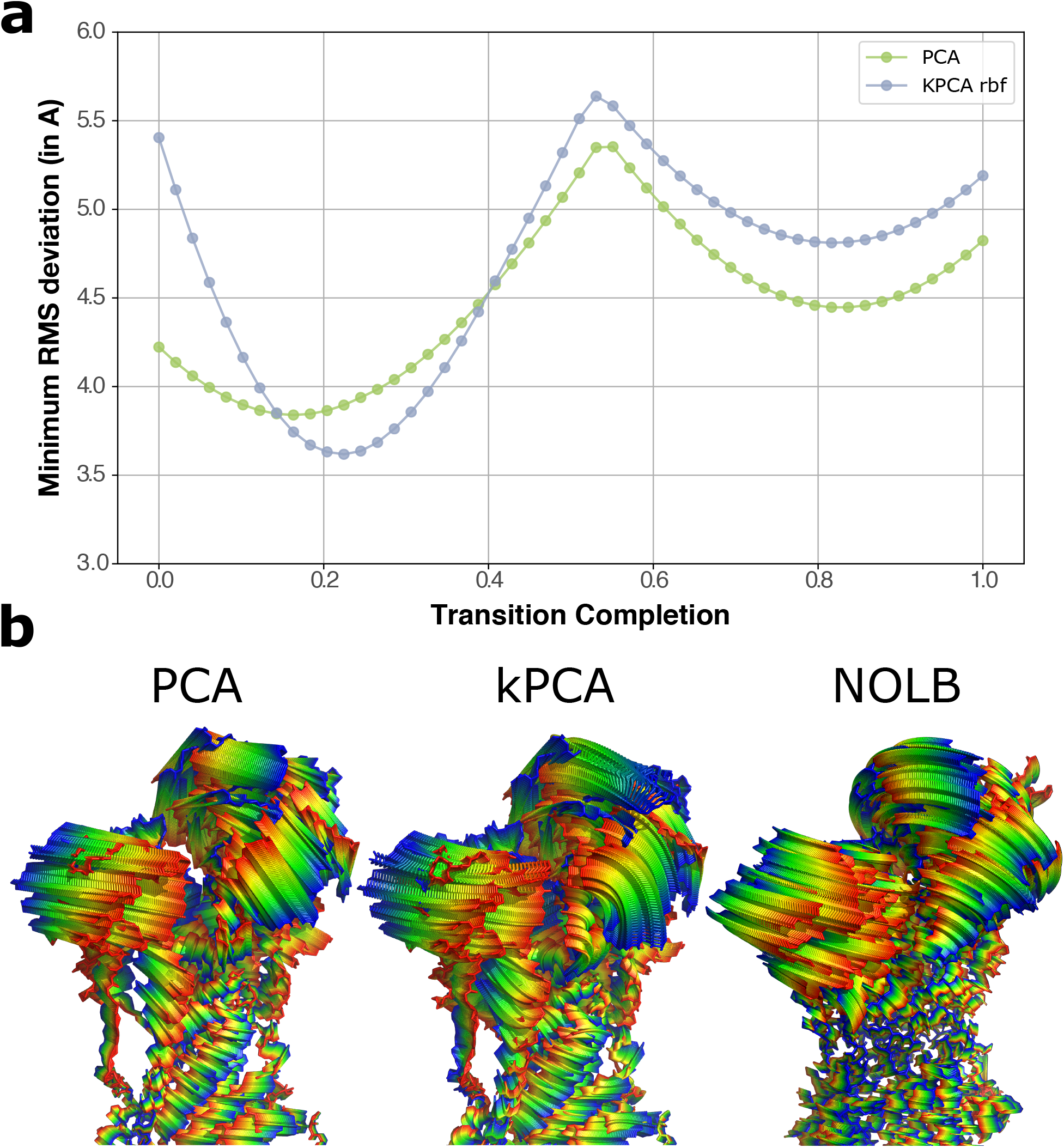
Interpolation trajectories for ATPase. The interpolation trajectories were computed between clusters 0 and 4, as depicted on Figure S9. For the kPCA, we used the RBF kernel with hyperparameters σ = 130 and α = 1 × 10^−10^. **a**. For each of the 50 conformations generated along the PCA and kPCA trajectories, we report its minimum RMS deviation (in Å) to the known experimental intermediate conformations from clusters 1, 2 and 3. **b**. The conformations generated by PCA (left) and kPCA (middle) are colored according to the transition completion, from blue to red. We compare them with the transition computed directly in the ambient space by NOLB between the conformations 1KJUA from cluster 0 and 1T5SA from cluster 4. NOLB extrapolates motions computed from instantaneous linear and angular velocities, defined with the normal mode analysis, to large amplitudes (*see Methods*).

### Influence of data uncertainty handling and reference conformation choice

We assessed the influence of accounting for uncertainty in the data by assigning a weight to each position proportional to the number of conformations where it was resolved (*see Methods*). In principle, this operation may impact the conformations’ superimposition and, as a consequence, their final coordinates, as well as the extracted motions. In practice, 95% of the ∼ 35 000 ensembles at 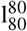 – excluding singletons and pairs, were not substantially altered by introducing position-wise uncertainty weights (**Supplementary Fig. S12**). They displayed the same displacement amplitude (±1 Å) and motion complexity (± 1 mode). When the weights were impactful, they effectively lowered the importance of large deviations in uncertain regions, *i*.*e*., poorly covered by the conformations, and prevented the associated motions, typically highly localised, from dominating the variance (**Supplementary Fig. S12**, red dots). Hence, the uncertainty weights tended to induce smaller deviations (Supplementary Fig. S12A), increased motion complexities (**Supplementary Fig. S12B**), and less dominant and more collective main motions (**Supplementary Fig. S12C-D**).

In addition, we performed two experiments probing the impact of choosing a different reference conformation. In the first one, we inverted the priority rules used to resolve ambiguities in the definition of the consensus sequence (see Methods). At a given position, in case of ambiguity, we would prefer a gap over an amino acid, thus favouring shorter reference conformations over longer ones, and a less frequent amino acid over a more frequent one, according to BLOSUM62 scores. Inverting the priority rules led to a different choice of reference in about 20% of the ∼35 000 collections. The displacement amplitude remained the same (±1 Å) in all cases and the motion complexity deviated by more than one mode in only one case (TrwK protein, from 6 to 4 modes). This analysis shows that changing the priority rules has a negligible impact on the results. In the second experiment, we applied a much more drastic change. Namely, we chose as alternative reference the conformation maximising the RMS deviation from the default reference. Moreover, we centred the data on the reference conformation, instead of the average conformation, prior to extracting the motions (see Methods). As expected, this setup yielded the most contrasted results, with about 57% of the ∼ 35 000 collections being impacted (**Supplementary Fig. S13**). It almost never happened that an ensemble consistently displayed a high motion complexity or a weakly contributing main motion for both references (**Supplementary Fig. S13B-C**). This result suggests that the ensembles exhibiting complex conformational rearrangements (*e*.*g*., loop deformations) among a bulk of conformations also include a few conformations comparatively far from all the others. The motions simplify when performing the PCA from the perspective of this minority. Normalising out the variance to focus on inter-residue correlations attenuates this effect (**Supplementary Fig. S14**).

## Discussion

This work proposes a new perspective on the variability of protein 3D conformations. It provides the community with conformational collections representing the multiple protein states available in the PDB and a fully automated versatile computational pipeline to build custom collections. In doing so, it contributes to the representation and managing of multiple conformational models of proteins. It enhances access and understanding of protein functional states and motions and facilitates predictive methods benchmarking. Both DANCE pipeline and the produced PDB-wide data are readily usable in other studies.

We chose to rely on classical principal component analysis because of its intuitive geometrical interpretation. It allows describing protein conformational variability with a limited set of orthogonal vectors interpretable as linear motions. By default, DANCE reports the number of PCA components required to explain 50%, 80%, 85%, 90%, 95%, and 99% of the total positional variance, thus providing a multi-resolution description of the complexity of the motions explaining the observed conformational diversity. We found that a few linear motions suffice to explain over 90% of the positional variance observed in the vast majority of the conformational collections. The high complexity exhibited by a few protein families may reflect nonlinear structural deformations or seemingly random fluctuations. For instance, protein kinases exhibit highly complex loop conformational rearrangements despite a well-conserved overall fold and only two metastable functional states. Our analysis helps to identify such cases to prioritise their in-depth characterisation with more sophisticated nonlinear dimensionality reduction techniques.

We designed DANCE for dealing primarily with single polypeptidic chains grouped based on sequence similarity. DANCE allows exploring different custom levels of sequence identity and coverage, thus providing a versatile framework for grouping the input 3D structures. Users who would like to save time may bypass the creation of the clusters and directly start from the pre-computed and weekly-updated clusters available through the RCSB PDB website. In addition, by default, DANCE analysis encompasses all polypeptidic chains found in the input 3D structures. These chains may be in different contexts and the motions extracted from the collections may be associated with the binding to a partner, as for BAX from the Bcl-2 family for instance. To ease interpretability, DANCE offers the users the possibility to restrict the context by excluding the protein chains engaged in oligomeric assemblies. Purely monomeric states represent about 15% of the ∼ 750K protein chains available from the PDB. Future improvements will include labelling complexes involving small molecules and accounting for them in the clustering. Furthermore, to go beyond sequence-based homology and the single-chain perspective, we have provided a proof-of-concept application study of DANCE’s usefulness for comprehensively describing continuous motions shared across very distant homologs comprising different numbers of chains. We showed that ABC proteins with a wide diversity of substrates and transport mechanisms share a highly collective high amplitude opening/closing motion underlying their functioning.

In addition, our work goes beyond a descriptive analysis by showing that classical manifold learning techniques can generate plausible conformations in the vicinity of the training set. These conformations could serve as starting points for further conformational exploration, e.g. with molecular dynamics simulations, or as targets in drug discovery campaigns. A potential strategy would be to give them as templates to RoseTTAFold All-Atom^92^ with a putative drug to guide the folding. The interpolation trajectories could provide insights into functional transitions involving substantial secondary structure rearrangements (e.g. membrane fusion proteins). The latter are particularly challenging to deal with for physics-based approaches, such as normal mode analysis^89^. Finally, our results can serve as baselines for evaluating more sophisticated approaches for predicting alternative conformations.

DANCE superimposes the conformations onto representative references and describes conformational variability as a set of linear motions of these references. This approach offers a multi-view perspective on a given collection of conformations, easing interpretability and allowing for augmenting data in a learning context. Nevertheless, radical differences between conformations, such as fold changes, might confound the superimposition. Another limitation comes from the dependency of the superimposition on the multiple sequence alignment heuristic. Ambiguities arising from sequence similarities might result in suboptimal 3D coordinates matching and, thus, in large deviations. Future improvements will explore multi-reference or reference-free probabilistic frameworks and more refined accounts of data uncertainty^93–97^.

## Data availability

We provide public access to the conformational collections compiled by DANCE from the PDB at two levels of sequence similarity, namely 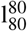 and 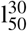 on Figshare^37^.This repository also contains the structural similarity-based ABC transporter conformational collection along with the supplementary **Movies S1** and **S2**. In addition, we provide detailed information about the benchmark set and the assessment of PCA and kPCA.

## Code availability

DANCE source codes are written in C/C++ and Python and are publicly available on GitHub at https://github.com/PhyloSofS-Team/DANCE. This repository also contains a Python wrapper allowing users to seamlessly run DANCE full pipeline. In addition, we provide example input 3D structures.

## Acknowledgements

We are grateful to Juliana Bernardes, Pablo Chacon, Tamas Hegedus, Anatoli Juditsky, and the Elixir 3D-Bioinfo Community members for insightful discussions and feedback. The Sorbonne Center for Artificial Intelligence (SCAI) provided a salary to VL and computational resources. This work has also been partially supported by the European Research Council under the European Union’s H2020 Framework Programme (2023–2028)/ ERC Grant agreement ID 101087830 awarded to EL.

## Author contributions statement

S.G. and E.L. designed research. V.L. and S.G. carried out the implementation. V.L., E.L. and S.G. produced and analysed the results. E.L. wrote the manuscript with support and feedback from all authors. S.G. and E.L. supervised the project.

