## Supplementary material for "Explaining Conformational Diversity in Protein Families through Molecular Motions"

### Supplemental Material for Explaining Conformational Diversity in Protein Families through Molecular Motions

#### Supplemental tables and figures

Supplemental Table S1: **Execution time on the PDB (748 297 protein chains)**

| **Step** | **Library or tool** | **(CPU)** | **Time#** |
|---|---|---|---|
| a- Sequence extraction | GEMMI | 16 | 9min 30s |
| b- Sequence clustering | MMseqs2 | 16 | 15s |
| c- Sequence alignment | MAFFT | 16 | 23min 54s |
| de- Structure extraction and alignment | GEMMI | 16 | 88min 2s |
| f- Linear motion extraction | NumPy | 16 | 11min 46s |
| **Total** | | | 2h 14min |

Supplemental Table S2: **Properties of the ensembles in the most conservative and the most relaxed set ups.**

| Property | Id., Cov. | Min. | 1st Quart. | Median | 3rd Quart. | Max. | Mean ± Sd |
|---|---|---|---|---|---|---|---|
| Ensemble size | 80,80 | 2.00 | 2.00 | 4.00 | 8.00 | 3 334 | 12.00 ± 45.80 |
| (in conformation) | 30,50 | 2.00 | 3.00 | 5.00 | 14.00 | 12694 | 24.64 ± 137.35 |
| Reference length | 80,80 | 5.00 | 112.00 | 207.00 | 335.00 | 4579.00 | 252.03 ± 223.35 |
| (in residue) | 30,50 | 5.00 | 81.00 | 165.00 | 304.00 | 4516.00 | 224.07 ± 231.74 |
| Max deviation | 80,80 | 0.00 | 0.52 | 1.10 | 2.45 | 72.93 | 2.39 ± 3.95 |
| (in Å) | 30,50 | 0.00 | 0.78 | 2.15 | 5.11 | 114.50 | 4.25 ± 5.88 |
| Motion complexity[a] | 80,80 | 1.00 | 2.00 | 3.00 | 4.00 | 118.00 | 3.89 ± 4.43 |
| (in mode) | 30,50 | 1.00 | 2.00 | 3.00 | 4.00 | 105.00 | 3.98 ± 4.45 |
| 1st mode contribution[a] | 80,80 | 7.30 | 50.00 | 64.90 | 80.90 | 100 | 64.85 ± 19.99 |
| (in percentage) | 30,50 | 7.30 | 49.90 | 65.40 | 82.20 | 100 | 65.32 ± 20.31 |
| 1st mode collectivity[a] | 80,80 | 0.30 | 13.30 | 30.20 | 50.90 | 98.20 | 33.07 ± 21.92 |
| (in percentage) | 30,50 | 0.30 | 15.90 | 29.40 | 48.90 | 98.20 | 33.26 ± 20.90 |

[a] To compute motion properties, we focused on the subset of ensembles with at least three members. Indeed, pairs of conformations trivially exhibit single-mode motions and are thus disregarded variability exhibited by pairs of conformations can be trivially explained by only one mode

Supplemental Table S3: **Properties of the ensembles chosen for benchmarking manifold learning techniques.**

| **Reference PDB Id** | **Protein name** | **Size** (# conf.) | **Length** (# res.) | **Mean deviation** (Å) | **Max deviation** (Å) | **Motion complexity** (# mode) | **1st mode contrib.** (%) | **1st mode coll.** (%) |
|---|---|---|---|---|---|---|---|---|
| 1AKEA | ADK | 42 | 214 | 2.48 ± 2.78 | 7.30 | 1 | 0.963 | 0.462 |
| 2WJPA | MurD | 22 | 435 | 2.58 ± 4.09 | 16.07 | 2 | 0.816 | 0.616 |
| 1IWOA | ATPase | 104 | 994 | 7.12 ± 4.19 | 15.84 | 3 | 0.597 | 0.481 |
| 5KOYB | ABC | 61 | 1181 | 5.95 ± 3.20 | 14.94 | 4 | 0.786 | 0.681 |
| 2KLRA | Crys | 23 | 82 | 2.08 ± 1.24 | 5.37 | 6 | 0.502 | 0.311 |
| 1AH6A | HSP90 | 52 | 213 | 1.13 ± 1.10 | 4.56 | 7 | 0.410 | 0.105 |
| 1NIWA | CALM | 388 | 136 | 10.38 ± 4.23 | 23.68 | 8 | 0.415 | 0.580 |
| 2G1TA | KIN | 122 | 271 | 2.85 ± 1.88 | 8.89 | 9 | 0.669 | 0.086 |
| 6YXWC | RAS | 744 | 167 | 1.56 ± 1.02 | 8.58 | 24 | 0.347 | 0.173 |
| 3J345 | CAP | 3334 | 231 | 4.06 ± 1.68 | 17.55 | 30 | 0.232 | 0.070 |

Supplemental Table S4: **Proportion of conformations reconstructed with high accuracy.**

| Reference | Protein | PCA (in %) | Poly-kPCA (in %) | RBF-kPCA (in %) | Sigmoid-kPCA (in %) | UMAP (in %) |
|---|---|---|---|---|---|---|
| 1AKEA | ADK | 83 | 83 | 83 | 83 | - |
| 2WJPA | MurD | 5 | 82 | 82 | 82 | 0 |
| 1IWOA | ATPase | 0 | 0 | 0 | 0 | 0 |
| 5KOYB | ABC | 18 | 28 | 30 | 18 | 0 |
| 2KLRA | Crys | 96 | 96 | 96 | 91 | 83 |
| 1AH6A | HSP90 | 92 | 90 | 90 | 92 | 90 |
| 1NIWA | CALM | 0 | 1 | 2 | 0 | - |
| 2G1TA | KIN | 93 | 93 | 93 | 93 | 57 |
| 6YXWC | RAS | 99 | 99 | 98 | 99 | - |
| 3J345 | CAP | 99 | 99 | 99 | 99 | - |

We consider reconstructions with RMSD errors lower than 2 Å as highly accurate. For the kPCA, we set the hyperparameters to the values leading to the lowest average error over each conformational collection (see **Supplementary Table S5**).

Supplemental Table S5: **Average reconstruction errors for unseen conformations and hyperparameter values.**

| Protein name | Method | Kernel type | sigma | alpha | $n_{neigh}$ | $d_{min}$ | Mean RMSD (Å) |
|---|---|---|---|---|---|---|---|
| ADK | pca | | | | | | 1.635 |
| ADK | kpca | rbf | 5.96e+03 | 1.00E-14 | | | 1.633 |
| ADK | kpca | poly | 13.3 | 2.81e+03 | | | 1.598 |
| ADK | kpca | sigmoid | 0.309 | 32.4 | | | 1.525 |
| MurD | pca | | | | | | 3.115 |
| MurD | umap | | | | 2 | 0.223 | 3.928 |
| MurD | kpca | rbf | 0.494 | 1.00E+05 | | | 1.774 |
| MurD | kpca | poly | 33.9 | 2.81e+03 | | | 1.755 |
| MurD | kpca | sigmoid | 1.00E+05 | 1.00E+05 | | | 1.774 |
| ATPase | pca | | | | | | 3.607 |
| ATPase | umap | | | | 9 | 0.112 | 5.161 |
| ATPase | kpca | rbf | 569 | 1.46e-13 | | | 3.567 |
| ATPase | kpca | poly | 910 | 5.96e-14 | | | 3.591 |
| ATPase | kpca | sigmoid | 2.33e+03 | 5.96e-14 | | | 3.606 |
| ABC | pca | | | | | | 2.599 |
| ABC | umap | | | | 2 | 1 | 3.911 |
| ABC | kpca | rbf | 569 | 5.96e-14 | | | 2.554 |
| ABC | kpca | poly | 569 | 3.56e-13 | | | 2.533 |
| ABC | kpca | sigmoid | 3.73e+03 | 5.96e-14 | | | 2.600 |
| Crys | pca | | | | | | 1.360 |
| Crys | umap | | | | 2 | 0.001 | 1.676 |
| Crys | kpca | rbf | 86.9 | 1.26e-11 | | | 1.356 |
| Crys | kpca | poly | 112 | 5.18e-12 | | | 1.357 |
| Crys | kpca | sigmoid | 0.193 | 5.43 | | | 1.302 |
| HSP90 | pca | | | | | | 0.719 |
| HSP90 | umap | | | | 4 | 0.889 | 0.865 |
| HSP90 | kpca | rbf | 54.3 | 7.54e-11 | | | 0.704 |
| HSP90 | kpca | poly | 222 | 2.44e-14 | | | 0.711 |
| HSP90 | kpca | sigmoid | 1.00E+03 | 1.00E-14 | | | 0.718 |
| CALM | pca | | | | | | 4.075 |
| CALM | kpca | rbf | 356 | 1.46e-13 | | | 3.525 |
| CALM | kpca | poly | 356 | 8.69e-13 | | | 3.531 |
| CALM | kpca | sigmoid | 356 | 1.6e-08 | | | 4.060 |
| KIN | pca | | | | | | 1.228 |
| KIN | umap | | | | 11 | 0.001 | 1.871 |
| KIN | kpca | rbf | 222 | 7.54e-11 | | | 1.225 |
| KIN | kpca | poly | 222 | 4.5e-10 | | | 1.226 |
| KIN | kpca | sigmoid | 910 | 5.96e-14 | | | 1.228 |
| RAS | pca | | | | | | 0.612 |
| RAS | kpca | rbf | 139 | 2.12e-12 | | | 0.605 |
| RAS | kpca | poly | 139 | 3.09e-11 | | | 0.606 |
| RAS | kpca | sigmoid | 910 | 2.12e-12 | | | 0.612 |
| CAP | pca | | | | | | 1.014 |
| CAP | kpca | rbf | 222 | 8.69e-13 | | | 0.986 |
| CAP | kpca | poly | 222 | 1.26e-11 | | | 0.989 |
| CAP | kpca | sigmoid | 356 | 4.5e-10 | | | 1.014 |

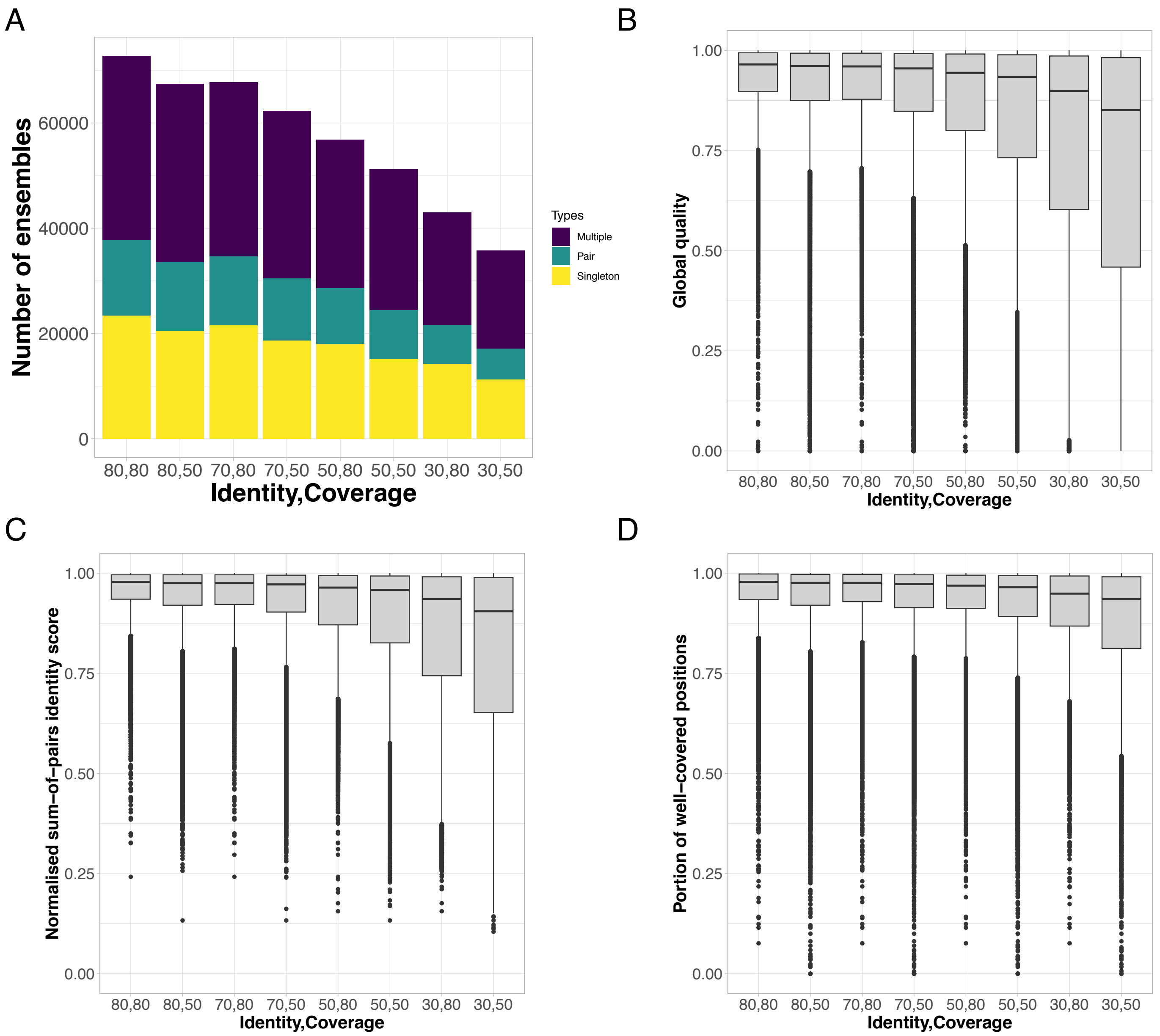


Supplemental Figure S1: **Global properties of the ensembles and their sequence alignments.** We report values computed across eight versions of the database, corresponding to eight combinations of sequence similarity and coverage thresholds. These combinations are given in x-axis. **A.** Number of singletons, pairs, and ensembles with at least 3 members. **B.** Distributions of sequence identity measured as a normalised sum-of-pairs scores with null mismatch and gap penalties. **C.** Distribution of coverage expressed as the fraction of positions with less than 80% gaps. **D.** Distribution of global alignment quality computed as a normalised sum-of-pairs scores with the following parameters: $\sigma_{match}$ = 1, $\sigma_{mismatch} = \sigma_{gap}$ = -0.5 (see *Methods*).

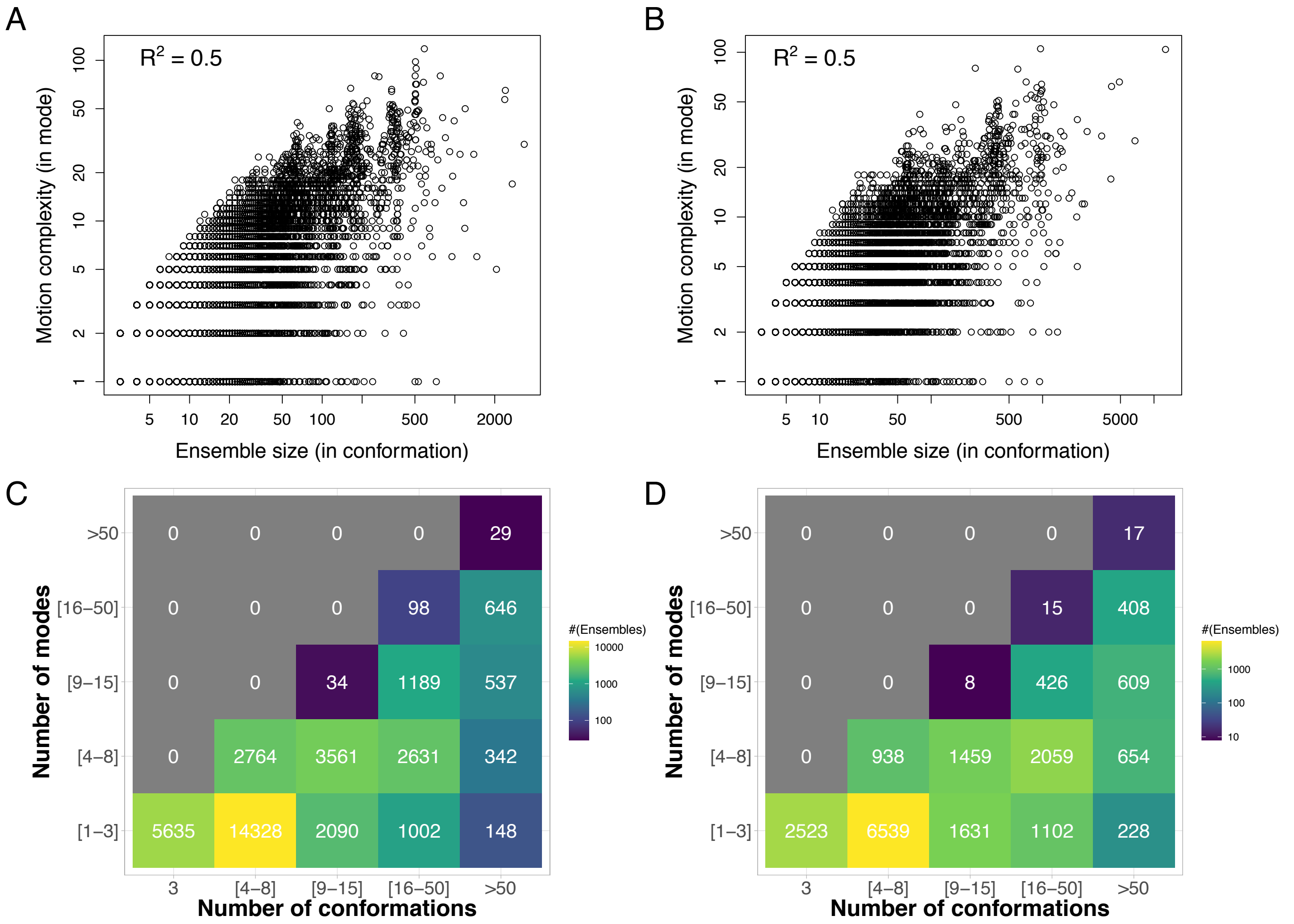


Supplemental Figure S2: **Influence of ensemble size on motion complexity** We report motion complexity, measured as the number of principal components or modes required to explain 80% of the positional variance, in function of the ensemble size, *i.e.* number of conformations. **A-B.** Scatterplots in log scale. **C-D.** Discretized heatmaps. We consider the most stringent set up, namely $l_{80}^{80}$ (A,C), and the most relaxed one, namely $l_{50}^{30}$ (B,D).

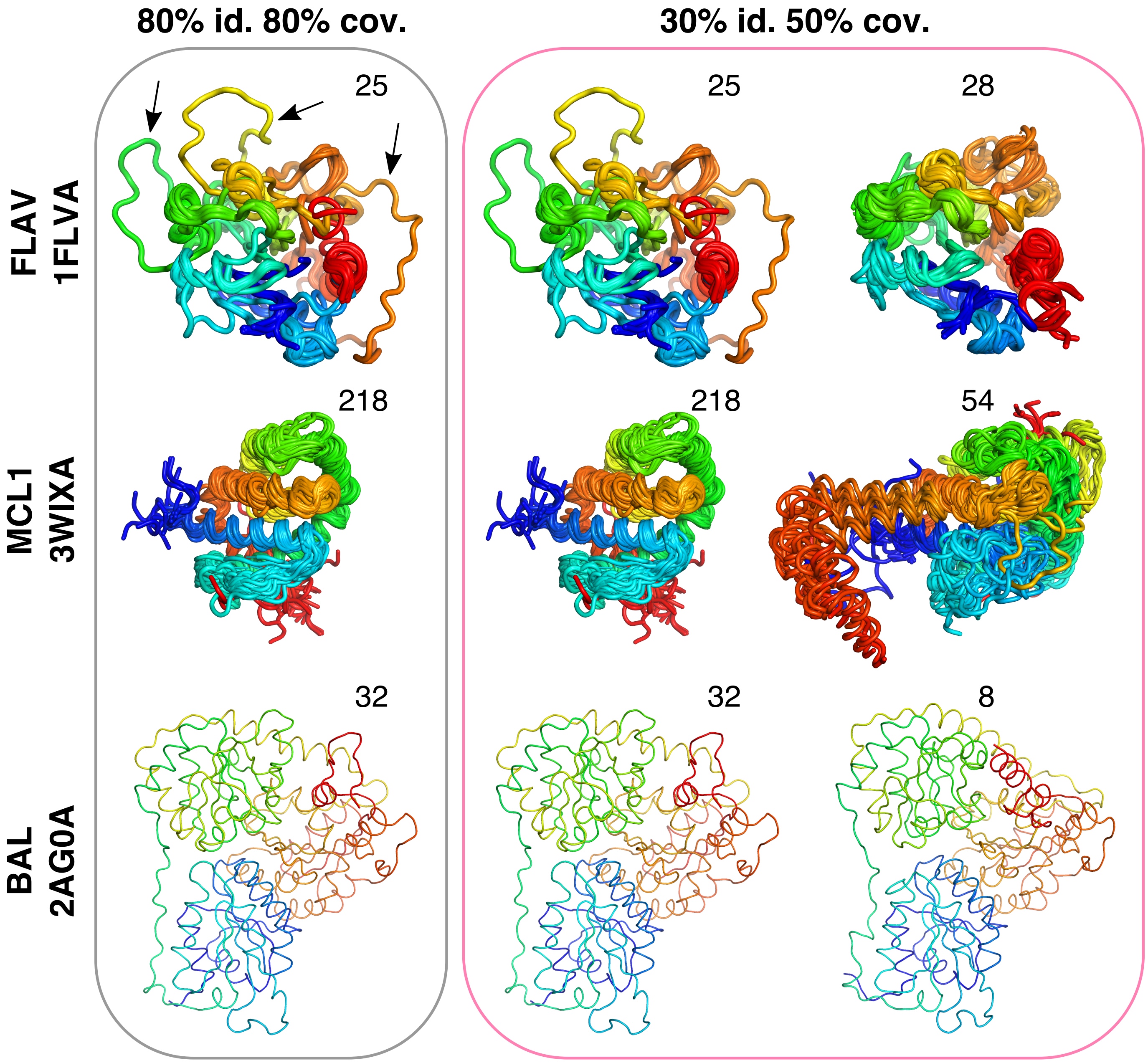


Supplemental Figure S3: **Expansion of three conformational ensembles upon relaxing sequence selection criteria.** We compare the set of conformations detected at two different levels of sequence similarity and coverage, namely $l_{80}^{80}$ (on the left) and $l_{50}^{30}$ (on the right). For the latter, we show separately the conformations already included in the ensemble at $l_{80}^{80}$ (on the left) and the new additional conformations (on the right). The number of conformations in each (sub)ensemble is given on top. The color code indicates the position in the sequence, from the N-terminus in blue to the C-terminus in red. The flavodoxin (FLAV) ensemble contains one partially unfolded conformation, highlighted with the arrows. Some properties of these three examples are reported in Figure 2.

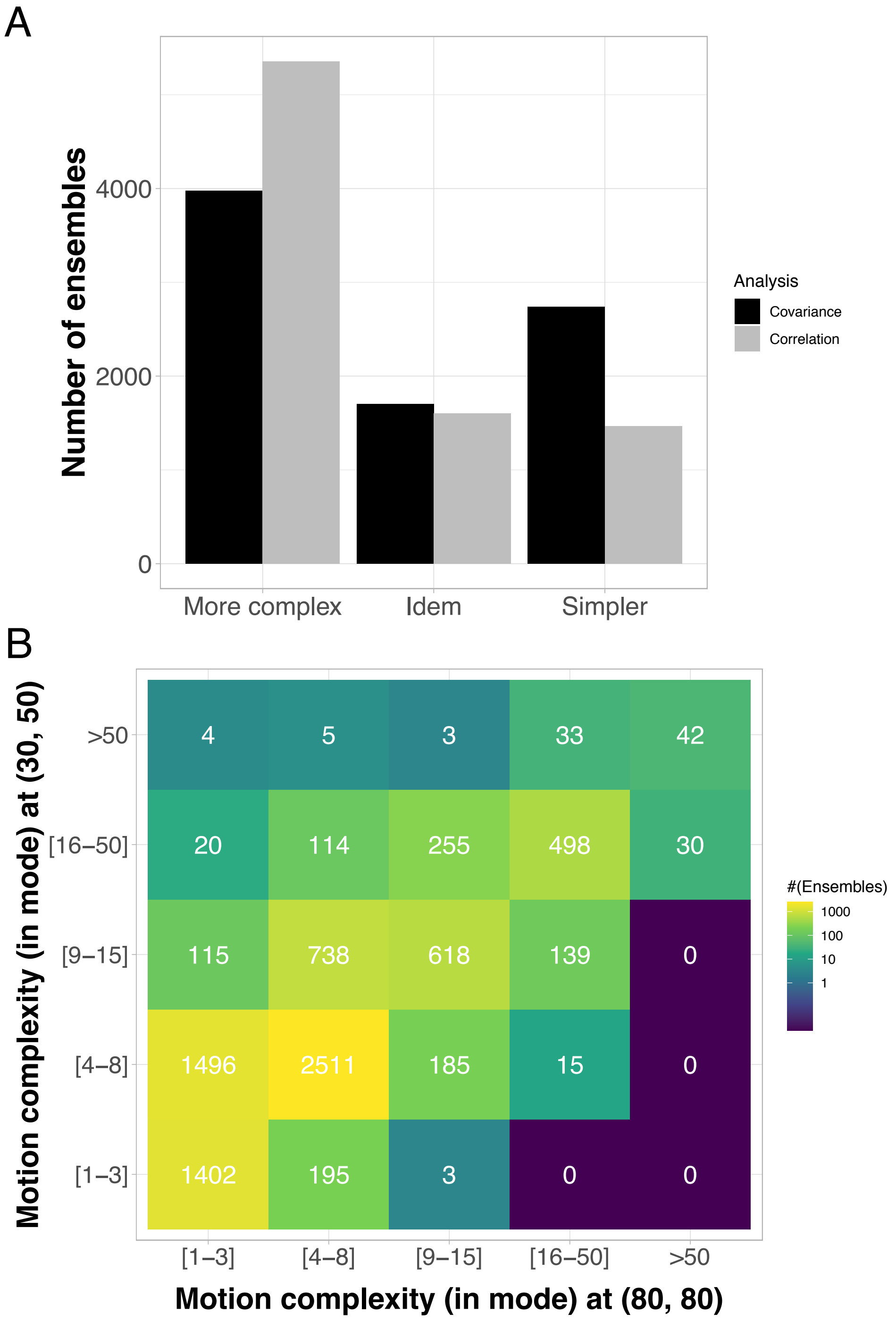


Supplemental Figure S4: **Evolution of motion complexity upon protein family expansion.** **A.** Number of ensembles where motion complexity increases, remains the same, or decreases between the most stringent and the most relaxed set ups. We extracted the motions from either the covariance (in black) or the correlation (in grey) matrix. **B.** Comparison of motion complexity estimated from the correlation matrix in the most stringent set up (x-axis) versus the most relaxed one (y-axis).

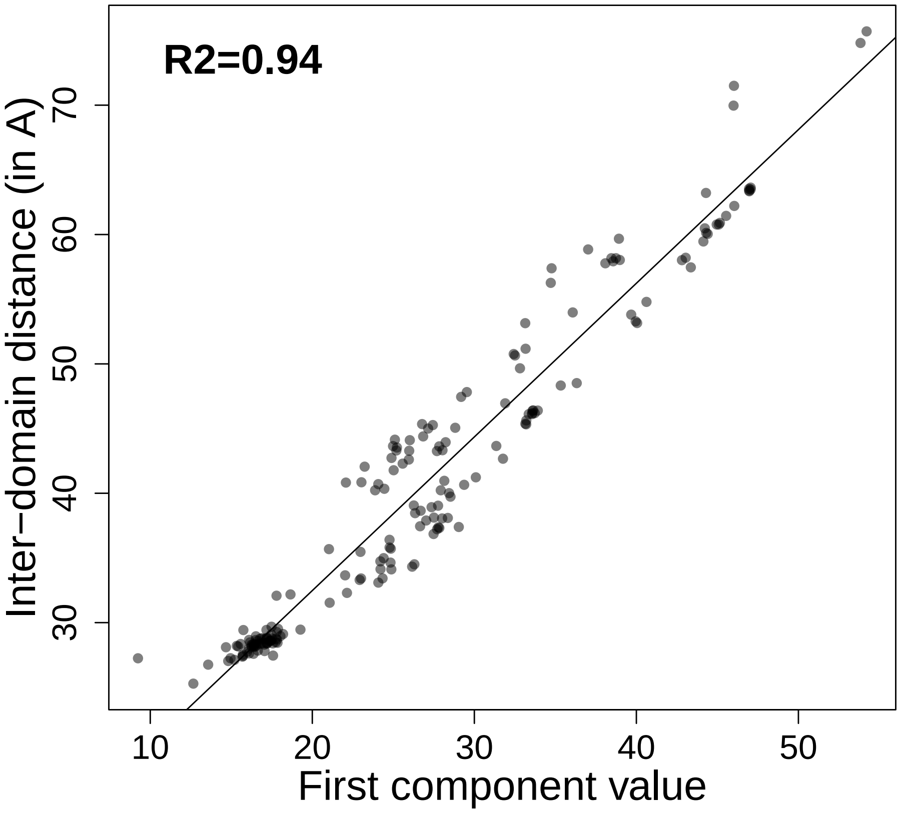


Supplemental Figure S5: **ABC protein opening in function of the first PCA component values.** The degree of opening of the ABC transporters is measured as the distance between the geometric centres of the two NBDs (in Å). The analysis is performed on the 188 conformations from the ABC structure similarity-based ensemble.

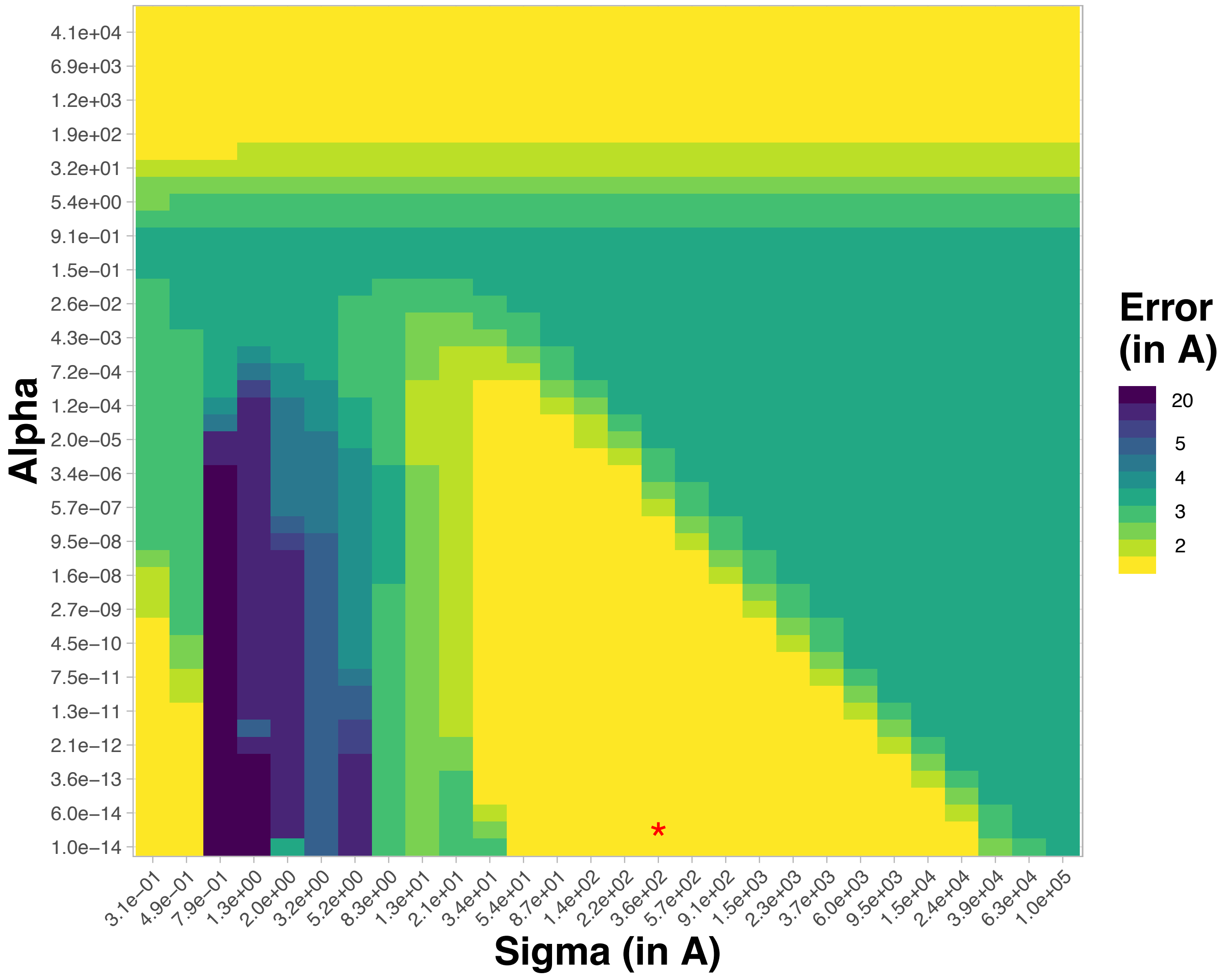


Supplemental Figure S6: **Systematic exploration of the two hyperparameters for kPCA-based conformation reconstruction.** We illustrate the influence of the hyper parameters $\sigma$ and $\alpha$ on the reconstruction error (in Å) for a randomly picked up conformation (4th one) from the ADK protein ensemble. The red star highlights the optimal parameter values. We used the RBF kernel.

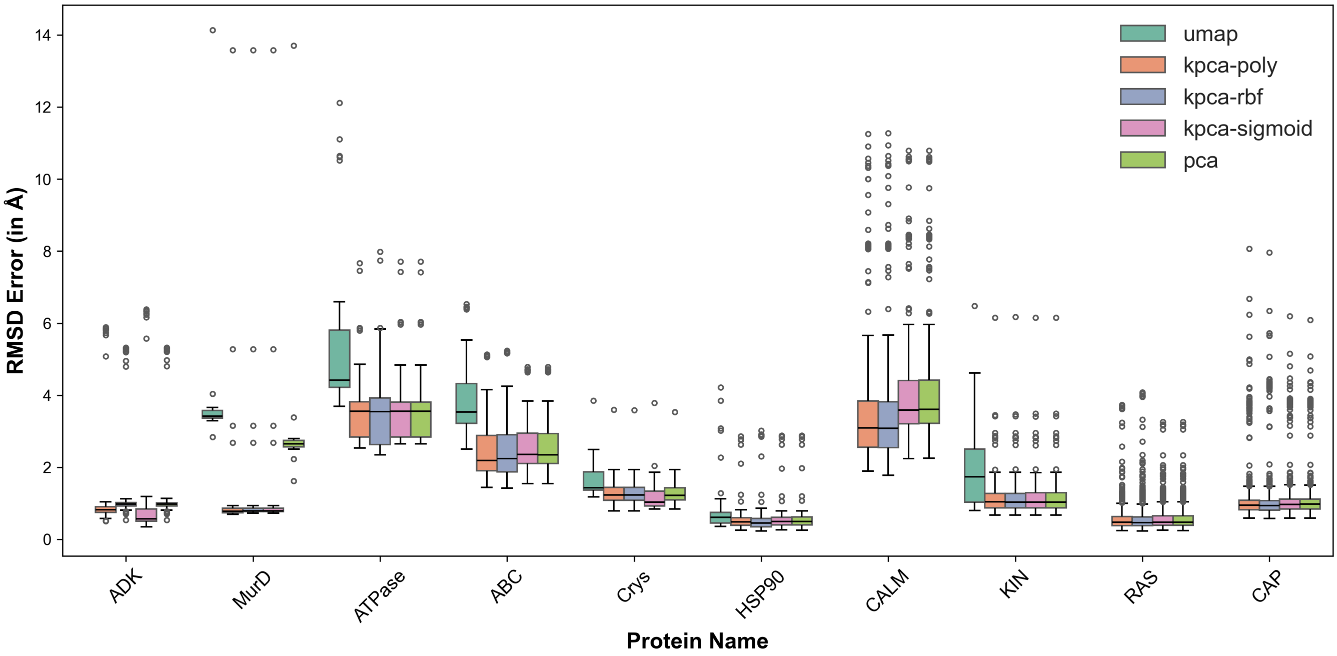


Supplemental Figure S7: **Distributions of the RMSD reconstruction errors (in Å) for each ensemble in the benchmark set.** We systematically reconstructed each conformation through a leave-one-cluster-out cross-validation procedure (see *Methods*). We set the hyperparameters of the kPCA and UMAP to the values yielding the best reconstruction, for each ensemble. The protein names in the x-axis are ordered according to motion complexity.

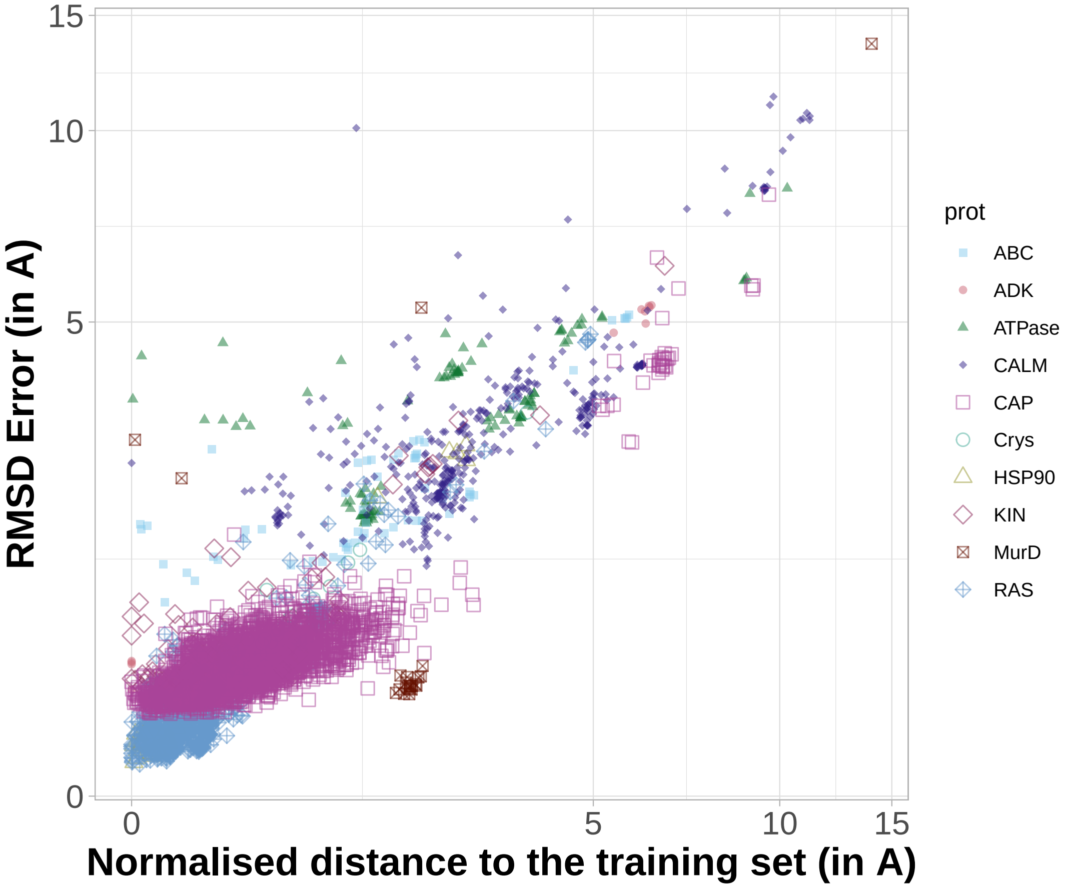


Supplemental Figure S8: **Reconstruction error in function of the distance to the training set for kPCA with RBF kernel.** The distance is computed between the test conformation and the convex hull defined by the training conformations in the low-dimensional representation space. It is normalised by the number of residues.

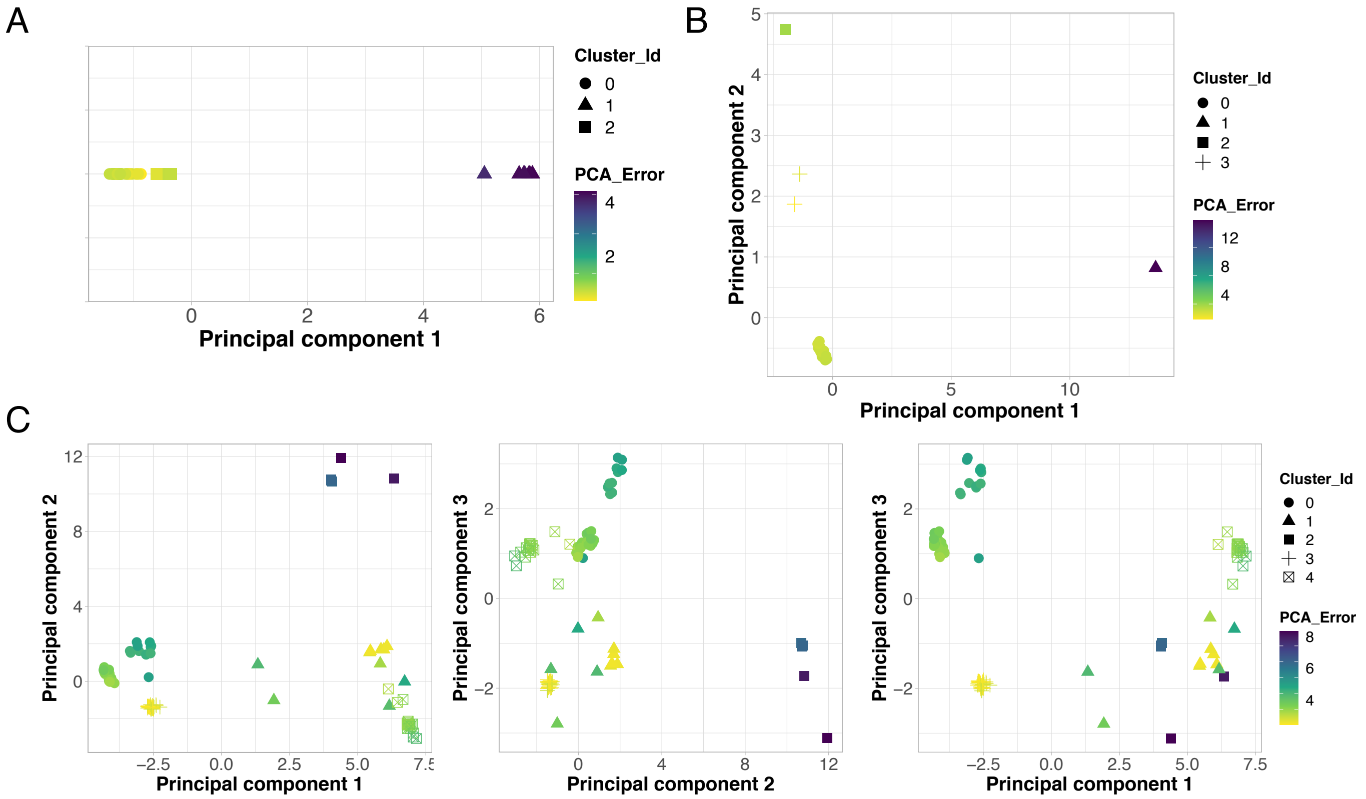


Supplemental Figure S9: **PCA feature spaces for three proteins from the benchmark.** We show the projections of the conformations in the $l$-dimensional PCA feature space, where $l$ is the number of principal components needed to explain 90% of the total positional variance, for ADK (A), MurD (B) and ATPase (C). The point shapes indicate the clusters to which the conformations belong as determined by k-means clustering where $k = l + 2$. The colors reflect the RMSD reconstruction error (in Å). We reconstructed each conformation using the principal components computed from the set of conformations not belonging to the same cluster.

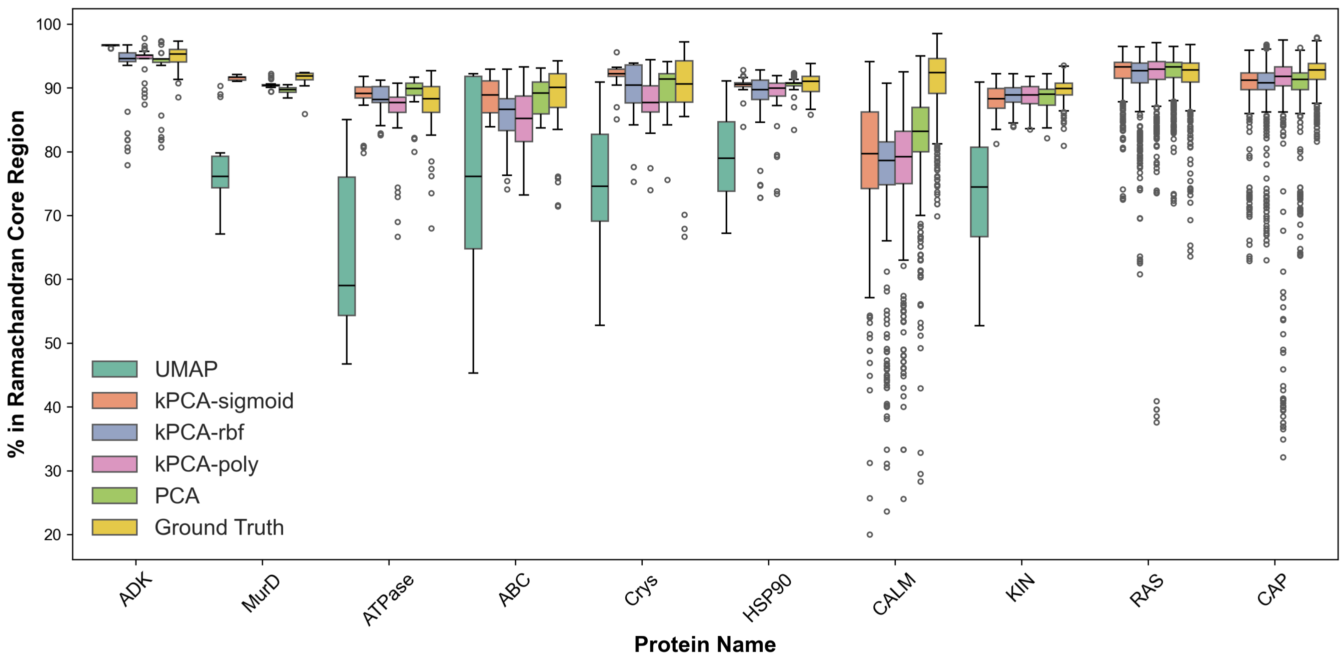


Supplemental Figure S10: **Distributions of the percentage of residues in the core region of the Ramachandran plot for each ensemble in the benchmark set.** We systematically reconstructed each conformation through a leave-one-cluster-out cross-validation procedure (see *Methods*). We set the hyperparameters of the kPCA and UMAP to the values yielding the best reconstruction, for each ensemble. The protein names on the x-axis are ordered according to motion complexity. The Ramachandran core region indicates the most favoured phi-psi angle combinations.

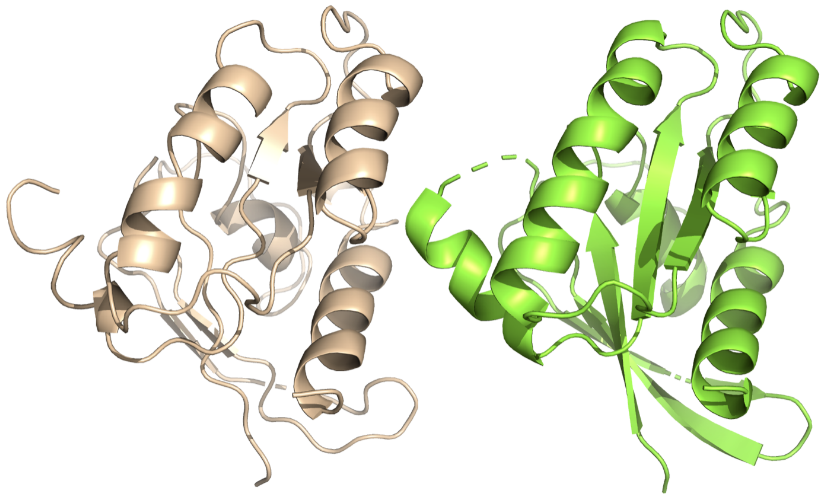

Supplemental Figure S11: **X-ray crystallographic structure of RAS (PDB code: 1PPL, chain A, in beige) and its PCA reconstruction (in green).** The PCA reconstruction displays a better secondary structure. The original structure has 63.9% of its residues in Ramachandran core region, while the PCA reconstruction has 96.4%.

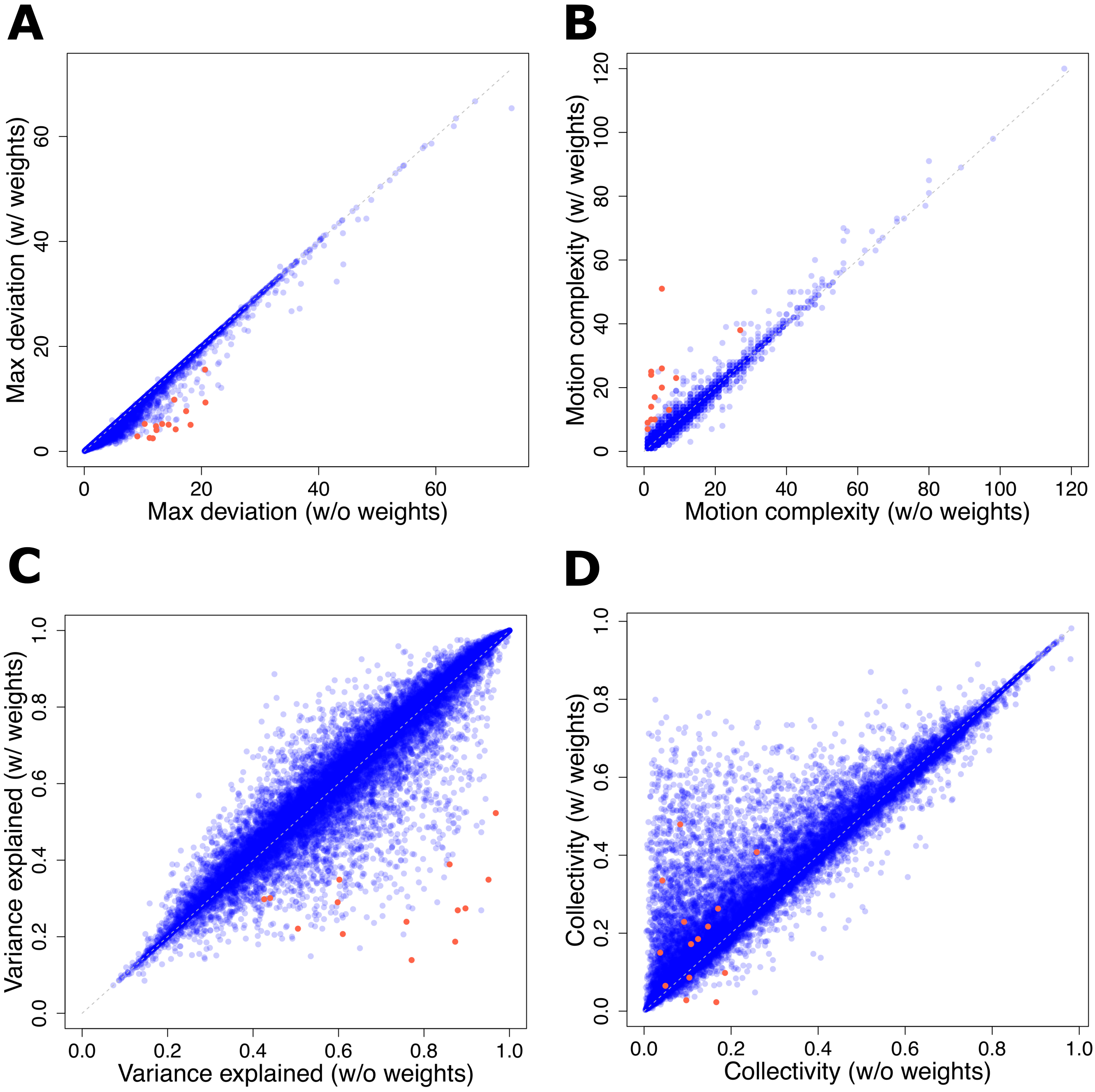


Supplemental Figure S12: **Influence of data uncertainty handling.** We compare some properties of the ensembles obtained at $l_{80}^{80}$ when applying the weighting scheme accounting for uncertainty versus without weights. We consider only the ensembles with at least 3 members. **A.** Largest deviation between any two conformations (in Å). **B.** Motion complexity (in mode). **C.** Percentage of the variance explained by the most contributing linear motion. **D.** Collectivity of the most contributing linear motion. We highlight the ensembles for which applying the weighting scheme leads to a maximum deviation decrease of more than 5Å and an increased motion complexity by more than 5 modes in red.

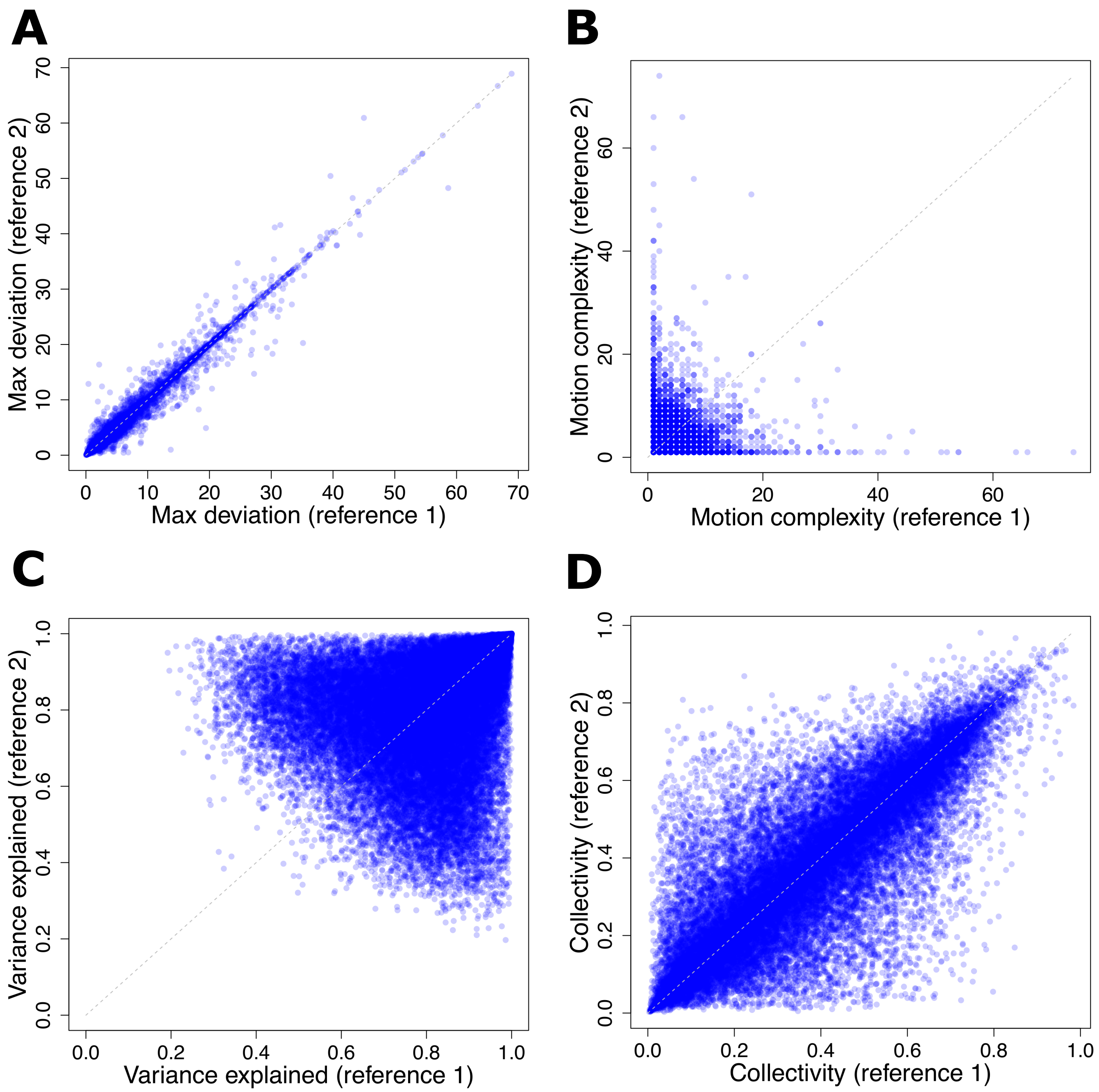


Supplemental Figure S13: **Influence of data conformation-specific centring.** We compare some properties of the ensembles obtained at $l_{80}^{80}$ when superimposing and centring the conformations with respect to two different references. The first one is the closest to the multiple sequence alignment consensus (see *Materials and methods*). The second one has the highest RMS deviation from the first one. We consider only the ensembles with at least 3 members. **A.** Largest deviation between any two conformations (in Å). **B.** Motion complexity (in mode). **C.** Percentage of the variance explained by the most contributing linear motion. **D.** Collectivity of the most contributing linear motion.

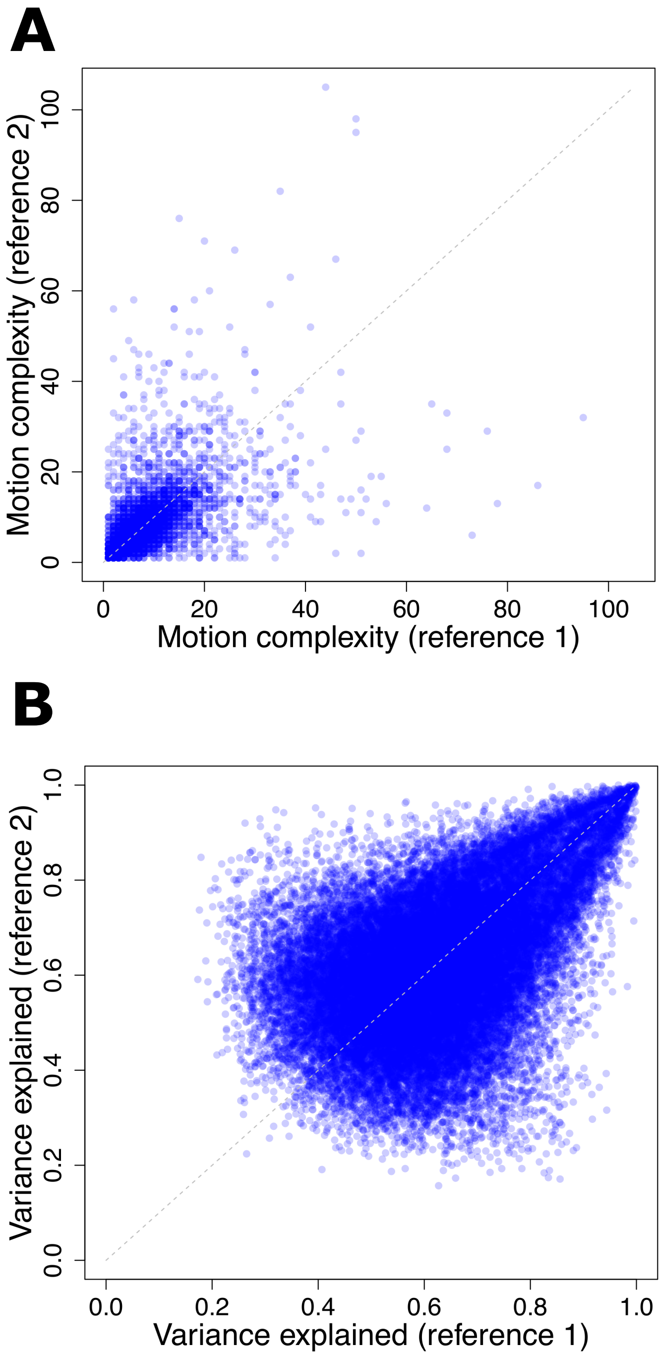


Supplemental Figure S14: **Influence of data conformation-specific centring when extracting motions from the correlation matrix.** We consider the ensembles with at least 3 members produced at $l_{80}^{80}$ when superimposing and centring the conformations with respect to two different references. The first one is the closest to the multiple sequence alignment consensus (see *Materials and methods*). The second one has the highest RMS deviation from the first one. **A.** Motion complexity (in mode). **B.** Percentage of the variance explained by the most contributing linear motion. These plots can be compared with panels B and C from Figure S9.
